# Targeting mitochondrial respiration and the BCL2 family in MYC-associated B-cell lymphoma

**DOI:** 10.1101/2020.11.22.390922

**Authors:** Giulio Donati, Micol Ravà, Marco Filipuzzi, Paola Nicoli, Laura Cassina, Alessandro Verrecchia, Mirko Doni, Simona Rodighiero, Federica Parodi, Alessandra Boletta, Christopher P. Vellano, Joseph R. Marszalek, Giulio F. Draetta, Bruno Amati

**Affiliations:** European Institute of Oncology (IEO) - IRCCS, Milan, Italy; IRCCS San Raffaele Scientific Institute, Milan, Italy; Translational Research to Advance Therapeutics and Innovation in Oncology (TRACTION), University of Texas MD Anderson Cancer Center, Houston, TX 77030, USA; Department of Genomic Medicine, University of Texas MD Anderson Cancer Center, Houston, TX 77030, USA

**Keywords:** Lymphoma, MYC, BCL2, OxPhos

## Abstract

Multiple molecular features, such as activation of specific oncogenes (e. g. *MYC*, *BCL2*) or a variety of gene expression signatures, have been associated with disease course in diffuse large B-cell lymphoma (DLBCL). Understanding the relationships between these features and their possible exploitation toward disease classification and therapy remains a major priority in the field. Here, we report that MYC activity in DLBCL is closely correlated with – and most likely a driver of – gene signatures related to Oxidative Phosphorylation (OxPhos). On this basis, we hypothesized that enzymes involved in Oxidative Phosphorylation, and in particular electron-transport chain (ETC) complexes, might constitute tractable therapeutic targets in MYC-associated lymphoma. Indeed, our data show that MYC sensitizes B-cells to IACS-010759, a selective inhibitor of ETC complex I. Mechanistically, IACS-010759 activates an ATF4-driven Integrated Stress Response (ISR), engaging the intrinsic apoptosis pathway through the transcription factor CHOP. In line with these findings, IACS-010759 shows synergy with the BCL2 inhibitor venetoclax against double-hit lymphoma (DHL), a high-grade form of DLBCL with concurrent activation of *MYC* and *BCL2*. Similarly, in BCL2-negative lymphoma cell lines, inhibition of the BCL2-related protein Mcl-1 potentiates killing by IACS-010759. Altogether, ETC complex I inhibition engages the ISR to lower the apoptotic threshold in MYC-driven lymphomas and, in combination with select BCL2-family inhibitors, provides a novel therapeutic principle against this aggressive DLBCL subset.

**Statement of significance:** This work points to OxPhos as a key MYC-activated process and a tractable therapeutic target toward personalized treatment of high-grade DLBCL, providing strong context-dependent cooperation with BH3-mimetic compounds.

## Introduction

Diffuse large B-cell lymphoma (DLBCL) is a heterogeneous disease with variable clinical course. While 50-60% of all patients achieve cure with standard immune-chemotherapy (R-CHOP, comprising rituximab, cyclophosphamide, doxorubicin, vincristine and prednisone), others succumb despite aggressive salvage regimens, including high-dose chemotherapy with autologous stem cell support, allogeneic transplantation or CAR T-cell therapy (1). Multiple genetic and molecular features have been associated with disease course in DLBCL, including translocation and/or over-expression of specific oncogenes (e. g. *MYC*, *BCL2*), mutational profiles, as well as transcriptome-based classifiers such as the so-called *cell-of-origin* (COO) and *comprehensive consensus clustering* (CCC) (1–5). While *MYC*, *BCL2* and COO-associated gene expression signatures are now commonly monitored in the clinic (1), other features have yet to make their way into routine practice, and full elucidation of their clinical relevance is still lacking. Moreover, complex combinatorial arrangements of the above features have far-reaching implications in disease classification and prognosis, some of which are just beginning to emerge (4–6).

A series of coincidental observations pointed to a hypothetical relationship between *MYC* and one of the signatures defined in the CCC model, termed “OxPhos” (henceforth CCC-OxPhos) based on its enrichment for genes involved in Oxidative Phosphorylation (2). First, *MYC* activation (whether assessed by gene rearrangement, protein expression or associated gene signatures) and CCC-OxPhos have both been linked to poor prognosis in DLBCL patients treated with R-CHOP (2,3,7–9), both behaving independently from the COO classification system (2–4). Second, MYC up-regulates multiple genes involved in mitochondrial transcription and translation (10–12) and increases the cells’ dependency upon those processes (11,12), both of which are required for assembly of the electron-transport chain (ETC) and oxidative phosphorylation. In line with this principle, tigecycline an antibiotic that inhibits the mitochondrial ribosome and impairs OxPhos activity (13,14) – showed increased toxicity toward either MYC-overexpressing cells (11,12) or DLBCL cell lines of the CCC-OxPhos subtype (15). Yet, whether MYC- and OxPhos-associated gene expression signatures might be related in DLBCL remains to be addressed: we show here that this is indeed the case, with high levels of correlation across multiple patient cohorts.

Altogether, the above observations led us to assess the potential of ETC inhibitors as therapeutic agents against MYC-associated high-grade DLBCL. We focused on IACS-010759 (16), a small-molecule inhibitor of ETC complex I that showed anti-tumoral activity in preclinical models of acute myeloid leukemia (AML), chronic lymphocytic leukemia, mantle cell lymphoma and lung cancer (16–19). Our data show that IACS-010759 preferentially induces apoptosis in MYC-overexpressing cells and cooperates with the BCL2 inhibitor venetoclax in killing *MYC/BCL2* DHL cells, as previously reported for Tigecycline (20). Furthermore, in BCL2-negative, *MYC*-translocated lymphoma cell lines, the cytotoxic activity of IACS was potentiated by the Mcl-1 inhibitor S63845. Hence, our data point to the possible use of ETC inhibitors in combination with distinct BH3-mimetic compounds (21) for the treatment of high-grade DLBCL, in a subtype- and patient-specific manner.

## Materials and Methods

### Analysis of DLBCL cohorts

For survival and correlation analyses in R-CHOP-treated DLBCL patients cohorts, we used gene expression data from six publicly available datasets (3–5,22–24). RNA-seq samples and patient’s survival data for Reddy et al. (3) and Ennishi et al. (22) where made available by the authors, and are accessible through the European Genome-phenome Archive (RRID: SCR_004944) at the European Bioinformatics Institute (https://www.ega-archive.org/; RRID: SCR_004727) with the accession numbers EGAS00001002606 and EGAS00001002657, respectively. RNA FastQ files were processed with the same pipeline used for our RNA-seq data, except for being aligned to hg19. Preprocessed data from Schmitz et al. (5) were downloaded from the Genomic Data Commons Data Portal (https://portal.gdc.cancer.gov/; RRID: SCR_014514) with project ID: NCICCR-DLBCL. The microarray data from Lenz et al. (23) (GSE10846), Sha et al. (24) (GSE117556) and Chapuy et al. (4) (GSE98588) were accessed through NCBI’s Gene Expression Omnibus. Probes were matched to the corresponding gene and, for those genes with more than 1 probe set, the mean gene expression was calculated. In all cases, data were first normalized to Transcripts Per kilobase Million (TPM) then, for each gene, the z-score across all samples was calculated. The Log-rank test was applied (survminer R package, https://rpkgs.datanovia.com/survminer/index.html, https://cran.r-project.org/web/packages/survival/index.html) for comparing survival curves of R-CHOP treated patients stratified according to the mean z-score of the expression of the genes belonging to the Hallmark MYC-V1 or OxPhos signatures. For the purpose of computing the linear correlation between signatures within a dataset, outliers were excluded from the calculation. Outliers were identified with the Inter-Quartiles Range (IQR) method as cases were the expression of the signature was not comprised within the following lower and upper boundaries: Lower boundary, 25^th^ quantile (IQR * 1.5); Upper boundary, 75^th^ quantile + (IQR * 1.5), where IQR = 75^th^ quantile – 25^th^ quantile. Moreover, in order to perform the analysis under stringent conditions, each pairwise correlation was calculated following exclusion of the genes that were shared between the two gene signatures (Supplemental Table 1).

**Table 1.** MYC- and OxPhos-associated gene signatures are correlated in DLBCL. Gene expression datasets from the DLBCL patient cohorts profiled in Chapuy et al. 2018 (4), Ennishi et al. 2019 (22), Lenz et al. 2008 (23), Reddy et al. 2017 (3), Schmitz et al. 2018 (5) and Sha et al. 2019 (24) were analyzed to define the pairwise correlations between each of the indicated MYC- and OxPhos-related gene expression signatures (Genes sets 1, 2), taken from the Hallmark collection (MYC-V1, MYC-V2 and Hallmark-OxPhos) and the CCC model (CCC-OxPhos). Within each cohort (indicated on the left), the rows represent each of the six pairwise comparisons (Gene set 1 and 2), ordered by decreasing Pearson correlation, with the indication of their statistical significance (p-value). Genes between shared between the compared signatures (Supplemental Table 1) were excluded from the calculation of each correlation.

| Cohort | Gene set 1 | Gene set 2 | Pearson Corr. | p-value |
| --- | --- | --- | --- | --- |
| Chapuy et al. 2018 | MYC-V1 | MYC-V2 | 0,841 | 8.08e-37 |
|  | Hallmark-OxPhos | MYC-V1 | 0,602 | 1.44e-14 |
|  | Hallmark-OxPhos | CCC-OxPhos | 0,595 | 3.55e-14 |
|  | Hallmark-OxPhos | MYC-V2 | 0,588 | 7.64e-14 |
|  | CCC-OxPhos | MYC-V1 | 0,471 | 9.67e-9 |
|  | CCC-OxPhos | MYC-V2 | 0,356 | 2.48e-5 |
| Ennishi et al. 2019 | MYC-V1 | MYC-V2 | 0,714 | 2.66e-47 |
|  | Hallmark-OxPhos | MYC-V1 | 0,618 | 1.01e-32 |
|  | Hallmark-OxPhos | MYC-V2 | 0,423 | 2.53e-14 |
|  | Hallmark-OxPhos | CCC-OxPhos | 0,297 | 1.89e-7 |
|  | CCC-OxPhos | MYC-V1 | 0,295 | 2.30e-7 |
|  | CCC-OxPhos | MYC-V2 | -0,007 | 0.90 |
| Lenz et al. 2008 | Hallmark-OxPhos | MYC-V1 | 0,888 | 1.43e-138 |
|  | Hallmark-OxPhos | CCC-OxPhos | 0,871 | 1.77e-126 |
|  | CCC-OxPhos | MYC-V1 | 0,819 | 2.07e-99 |
|  | MYC-V1 | MYC-V2 | 0,667 | 1.29e-53 |
|  | Hallmark-OxPhos | MYC-V2 | 0,573 | 8.31e-37 |
|  | CCC-OxPhos | MYC-V2 | 0,380 | 2.2.7e-15 |
| Reddy et al. 2017 | CCC-OxPhos | MYC-V1 | 0,744 | 6.45e-127 |
|  | Hallmark-OxPhos | MYC-V1 | 0,700 | 1.94e-106 |
|  | Hallmark-OxPhos | CCC-OxPhos | 0,517 | 4.27e-50 |
|  | MYC-V1 | MYC-V2 | 0,505 | 7.40e-47 |
|  | Hallmark-OxPhos | MYC-V2 | 0,503 | 3.45e-47 |
|  | CCC-OxPhos | MYC-V2 | 0,396 | 3.50e-28 |
| Schmitz et al. 2018 | MYC-V1 | MYC-V2 | 0,731 | 1.23e-73 |
|  | Hallmark-OxPhos | MYC-V1 | 0,686 | 1.58e-62 |
|  | Hallmark-OxPhos | CCC-OxPhos | 0,593 | 3.77e-43 |
|  | Hallmark-OxPhos | MYC-V2 | 0,472 | 9.39e-26 |
|  | CCC-OxPhos | MYC-V1 | 0,454 | 9.03e-24 |
|  | CCC-OxPhos | MYC-V2 | 0,115 | 0.02 |
| Sha et al. 2019 | CCC-OxPhos | MYC-V1 | 0,813 | 7.35e-206 |
|  | Hallmark-OxPhos | MYC-V1 | 0,715 | 1.12e-136 |
|  | MYC-V1 | MYC-V2 | 0,663 | 4.71e-109 |
|  | Hallmark-OxPhos | CCC-OxPhos | 0,602 | 1.35e-86 |
|  | Hallmark-OxPhos | MYC-V2 | 0,585 | 7.14e-81 |
|  | CCC-OxPhos | MYC-V2 | 0,557 | 5.11e-72 |

### Cell lines and Xenograft model

The human lymphoma cell lines SU-DHL-6, SU-DHL-4, DOHH-2, Karpas 422, OCI-LY7 and Ramos, the murine lymphoid precursor cell lines Ba/F3 and FL5.12, as well as their derivatives were grown as described in the Supplemental Materials and Methods section. FL5.12 cells were a gift from Pier Giuseppe Pelicci, and other cell lines were imported from the DSMZ or ATCC repositories (https://www.dsmz.de; https://www.lgcstandards-atcc.org); all lines were stocked and made available by IEO’s core Tissue Culture facility, where they were also tested for mycoplasma infection.

For the analysis of drug responses *in vivo*, 10^6^ SU-DHL-6 cells were transplanted subcutaneously in irradiated (3 Gray) CD1-nude nu/nu mice. Treatment started upon the appearance of measurable tumors. Tumor volumes were assessed from the start of the treatment every two days with a digital caliper and calculated as 1/2 length×width^2^ (mm^3^). The following treatment schemes were used: daily oral gavage with venetoclax and IACS-010759 (spaced by ~8 hours) for 5 days, followed by two days off and a repeat of the same scheme, for a total of 12 days. Venetoclax was dissolved in 60% Phosal 50 PG (Lipoid), 30% Polyethylene glycol (Aldrich), 10% Ethanol; IACS-010759 was suspended in 0.5% methylcellulose (Merck Life Science). During the experiment, two animals treated with the highest daily dose of IACS-010759 (15mg/kg), given alone and in combination venetoclax respectively, showed signs of toxicity, thus the treatment was discontinued and the animals were pulled out from the respective group.

Experiments involving animals were done in accordance with the Italian Laws (D.lgs. 26/2014), which enforces Dir. 2010/63/EU (Directive 2010/63/EU of the European Parliament and of the Council of 22 September 2010 on the protection of animals used for scientific purposes), and authorized by the Italian Health Ministry with project nr. 70/2019-PR.

### Oxygen consumption rate (OCR) and respiratory parameters

OCR was measures on the Seahorse XFe96 Analyzer (Agilent Technologies) using SeaHorse XF Cell Mito Stress Test, following the manufacturer’s instructions. Briefly, on the day of the assay, cells were counted and attached to 96-well Seahorse cell culture microplates, pre-coated with Corning™ Cell-Tak (Life Sciences) according to the manufacturer’s instructions, at a density of 80,000 cells/well. Cells were seeded in at least 8 wells per experimental condition, in XF RPMI Medium pH 7.4 with 1 mM HEPES (Agilent Technologies), supplemented with 2.75 mM glucose, 1 mM sodium pyruvate, 2 mM L-glutamine and, where specified, 100nM OHT, 135 nM IACS-010759, 100 nM venetoclax. The plates were incubated at 37 °C for 1 h in a non-CO2 incubator. After OCR baseline measurements, oligomycin A, Carbonyl cyanide 4-(trifluoromethoxy)phenylhydrazone (FCCP), and antimycin A/rotenone were added sequentially to each well, to final concentrations of 1 μM, 1.5 μM and 0.5 μM, respectively. Results were normalized by cell number, measured at the end of the experiment using the CyQUANT Cell Proliferation Assays (Thermo Fisher Scientific). Data are expressed as pmol of oxygen per minute per arbitrary units (pmol/min/a.u.). Respiratory parameters were calculated with the Wave Desktop 2.6 software and shown as mean ±SEM of 3 independent experiments.

Additional experimental procedures, measurements and analytical tools are described in detail in the Supplemental Materials and Methods section.

## Results

### MYC- and OxPhos-associated gene expression signatures are correlated in DLBCL

To address a possible relationship between MYC and OxPhos in DLBCL, we determined the pairwise correlations between four reference signatures, comprising either MYC-regulated genes (Hallmark-MYC-V1 and -MYC-V2) or genes related to oxidative phosphorylation (Hallmark-OxPhos and CCC-OxPhos), in six independent DLBCL datasets (3–5,22–24). Remarkably, the MYC- and OxPhos-associated signatures, in particular MYC-V1 and Hallmark-OxPhos, showed significant positive correlation in all patient cohorts (Table 1, supplemental Figure 1A). Most noteworthy, those two signatures also behaved very similarly as predictive biomarkers in R-CHOP-treated patients, showing significant association with worse outcome in two of the cohorts, loose association in one, and none in three others (supplemental Figure 1B, C). Thus, while variable in terms of their predictive value in R-CHOP-treated patients, the MYC- and OxPhos-associated signatures were highly correlated in all DLBCL cohorts.

**Figure 1.**
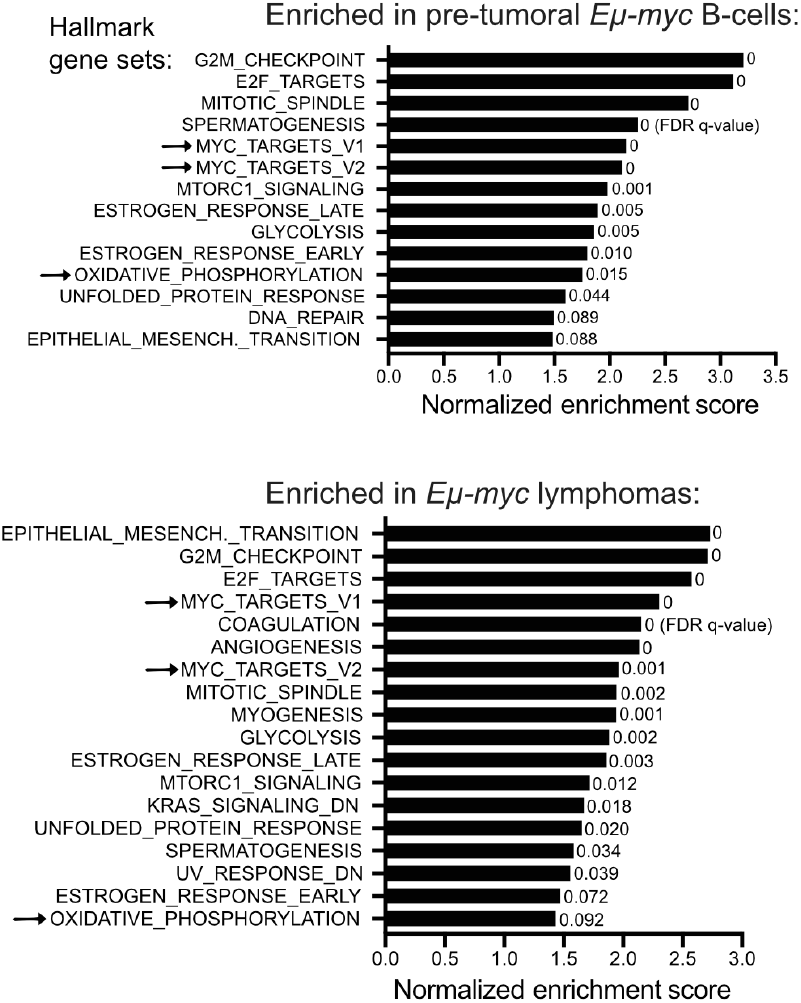
Biological processes enriched during Eμ-Myc driven lymphomagenesis. Our previous RNA-seq data (25) were used to address the enrichment of gene sets from the Hallmark collection and the CCC model (2) in pre-tumoral Eμ-*myc* B-cells (top) and lymphomas (bottom), relative to control non-transgenic B-cells. The plot shows the enriched genes signatures (FDR q-value < 0.1), ranked according to their normalized enrichment score. Only a subset of the Hallmark-associated biological pathways, but none of the CCC signatures, reached this threshold. The MYC target and OxPhos gene signatures are marked by an arrow.

We then addressed the behavior of the Hallmark and CCC-derived gene signatures during MYC-induced lymphomagenesis, based on previous RNA-seq data in Eμ-*myc* transgenic mice (25). Relative to control non-transgenic B-cells, either pre-tumoral Eμ-*myc* B-cells or late-stage lymphomas enriched not only for the MYC signatures, as expected, but also for Hallmark-Oxphos (albeit with lower significance in the lymphomas, owing most likely to their clonal heterogeneity) (26), while CCC-Oxphos showed no significant enrichment (Figure 1).

Altogether, the above data point to OxPhos as one of the positively regulated gene programs – among others (Figure 1) (25) – in MYC-driven lymphomagenesis. We previously reported that the same is true for up-regulation of the mitochondrial translation machinery (11), which is itself required for OxPhos activity (13,14,20). We thus hypothesized that, like mitochondrial ribosomes (11,12,20), OxPhos activity might represent a tractable therapeutic target in MYC-associated lymphoma.

### MYC sensitizes B-cells to IACS-010759-induced killing through the intrinsic apoptotic pathway

In order to address whether enhanced Myc activity may increase the sensitivity to OxPhos disruption, we transduced two murine B-cell progenitor lines (FL5.12 and Ba/F3) with retroviral vectors driving constitutive expression of a 4-hydroxytamoxifen (OHT)-dependent MycER chimera (hereafter FL^MycER^ and BaF^MycER^). At the phenotypic level, 48 hours of OHT treatment had no noticeable impact on either cycle transit times or cell death (supplemental Figure 2A, B), owing most likely to already maximal division rates in the basal state (ca. 12 hours) and to the presence of survival factors in the culture medium (27). Yet, as expected, OHT treatment led to the induction of known MYC-activated mRNAs, as well as suppression of endogenous *Myc* mRNA and protein (supplemental Figure 2C, D). We then used RNA-seq to profile the response to OHT treatment in FL^MycER^ cells: as in other cell types (25), MycER activation caused both up- and down-regulation of discrete sets of differentially expressed genes (DEGs: ca. 1200 each; supplemental Figure 2E). At the computational level, this response showed two typical features of MYC-regulated transcription: first, a search for upstream regulators confirmed MYC as its main driver (supplemental Figure 2F); second, the Hallmark-MYC-V1 gene set was enriched among OHT-induced DEGs (supplemental Figure 2G). Altogether, FL^MycER^ and BaF^MycER^ cells provide a reliable model for the study of synthetic-lethal interactions between MYC and selected drugs.

**Figure 2.**
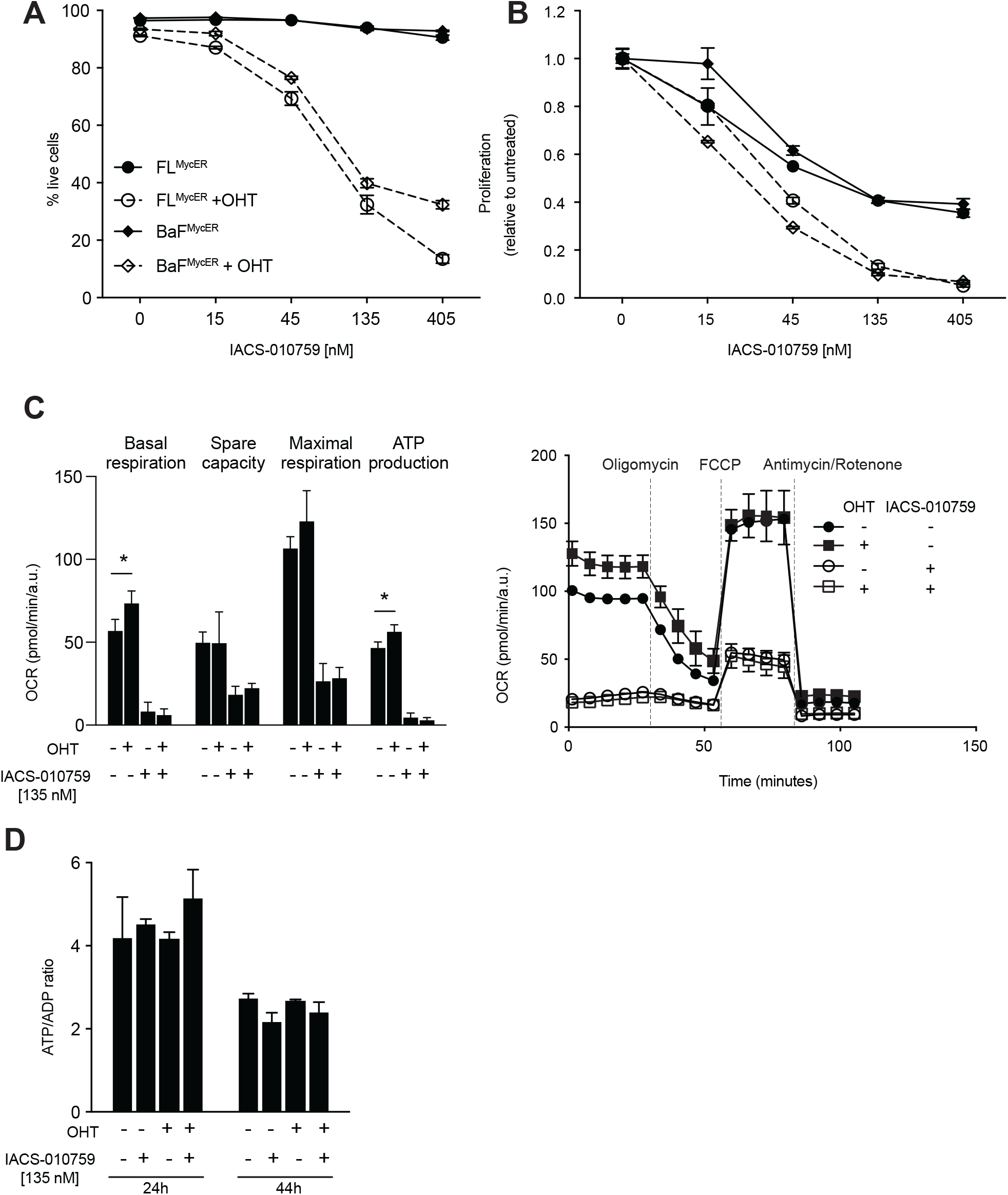
Elevated MYC activity sensitizes B-cells to IACS-010759. FL^MycER^ and BaF^MycER^ cell were pre-treated or not with OHT (100 nM, 48h), followed by 48 hours of IACS-010759 treatment at the indicated concentrations. **(A)** Percentage of live cells relative to total (live + dead) cells in each culture condition. **(B)** Proliferation index, based on viable cell counts at the end of the treatment, with each IACS-010759 treated sample normalized relative to its untreated control (either with, or without OHT). In (A) and (B), all OHT-primed samples treated with ≥ 45 nM IACS-010759 showed significant differences relative to their controls (p < 0.0001). Error bars in both panels: SD (n=3). Cell count and viability were determined by Propidium Iodide staining 48 hours after addition of IACS-010759 to the culture medium. **(C)** Left: basal respiration, spare capacity, maximal respiration, and respiration-coupled ATP production, averaged from 3 independent mitochondrial stress test profiles in FL^MycER^ cells, treated as indicated (Error bars: SEM; * p < 0.05); a representative profile is shown on the right. **(D)** ATP/ADP ratios in FL^MycER^ cells, treated as indicated. Error bars: SD (n=3).

We thus proceeded to treat FL^MycER^ and BaF^MycER^ cells with OHT and the ETC complex I inhibitor IACS-010759 (16). In the absence of OHT, increasing concentrations of IACS-010759 exerted no - or little cytotoxic activity; instead, pre-treatment with OHT markedly sensitized both cell lines to dose-dependent killing by IACS-010759 (Figure 2A). As seen in other cell types (16,28), this occurred with reduced amounts of glucose (2.75 mM) but not in standard high-glucose media (11 mM) (supplemental Figure 3A). Moreover, while killing depended upon MycER activation (Figure 2A), IACS-010759 was still cytostatic in the absence of OHT (Figure 2B). Finally, OHT promoted killing by another complex I inhibitor, rotenone (supplemental Figure 3B). Hence, MycER activation sensitized cells to inhibition of ETC complex I.

**Figure 3.**
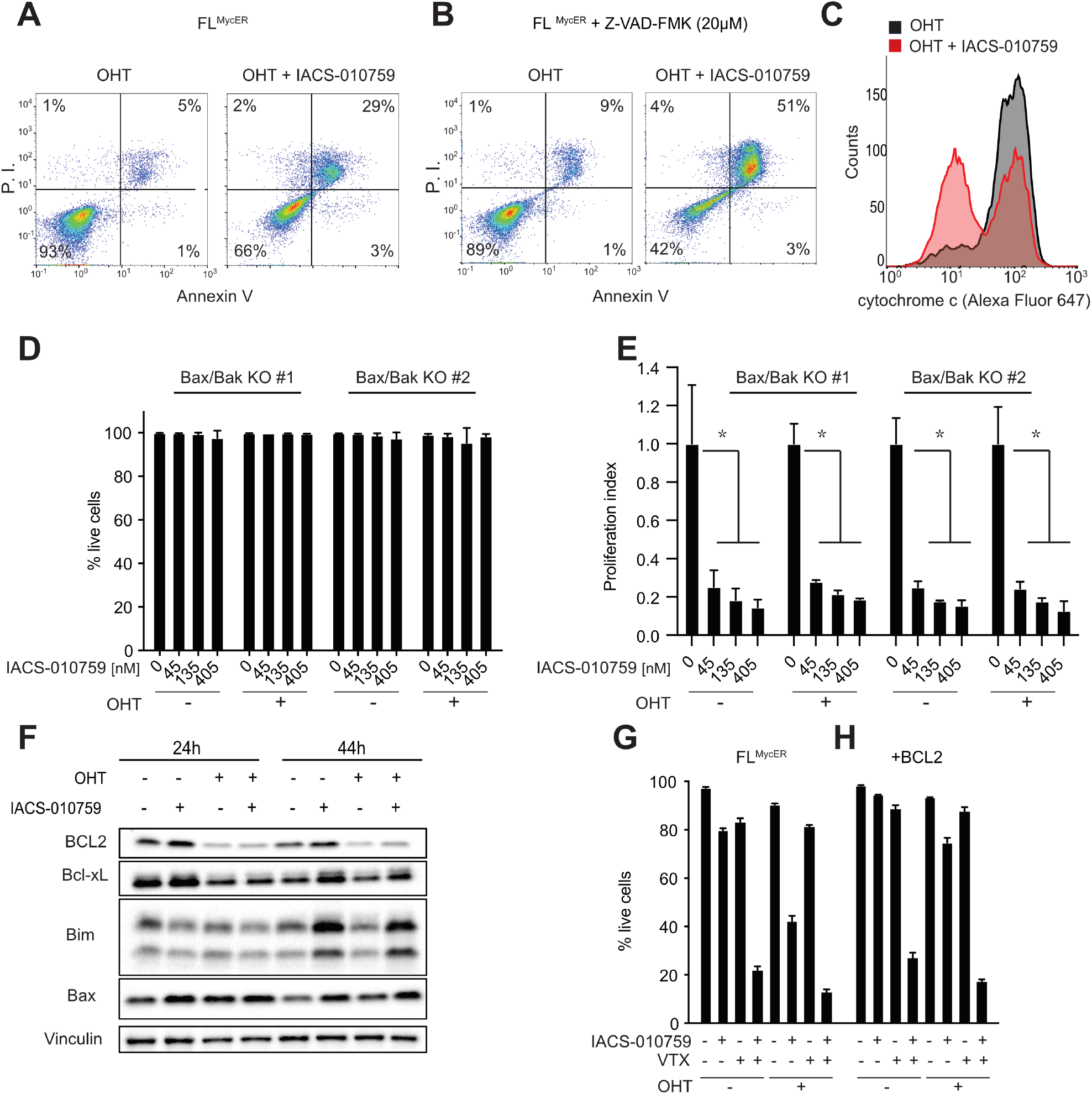
IACS-010759 activates intrinsic apoptosis in Myc overexpressing cells. FL^MycER^ cells were sequentially treated with OHT (100 nM, 48h), followed by IACS-010759 (135 nM, unless otherwise indicated; all 48h). (**A**) Apoptotic cell death was assayed by FACS analysis of Propidium Iodide (P.I.) and Annexin V staining (both shown as arbitrary fluorescence units). (**B**) As in (A) with the addition of 20 μM Z-VAD-FMK together with IACS-010759. (**C**) Mitochondrial retention of cytochrome c, evaluated by anti-cyt. c staining and FACS analysis of digitonin-permeabilized FL^MycER^ cells. (**D**) Percentage of live cells and (**E**) proliferation index (as defined in Figure 2) in two independent Bax/Bak double knockout FL^MycER^ clones pre-treated or not with OHT, then treated with IACS-010759 for 48 hours. Error bars: SD (n=3). *P < 0.001 vs. IACS-010759-untreated control. (**F**) Immunoblot analysis of the indicated BCL2-family members in FL^MycER^ cells pre-treated or not with OHT, followed by IACS-010759 at the indicated times. Vinculin was used as loading control. (**G, H**) Percentage of live cells in parental FL^MycER^ (G) and FL^MycER^ cells over-expressing BCL2 (H). The cells were pre-treated or not with OHT (100 nM, 48h), then treated for 48 hours with IACS-010759 (135 nM) and/or venetoclax (VTX; 100 nM), as indicated. Error bars: SD (n=3).

Further experiments in FL^MycER^ cells confirmed that the effect of IACS-010759 was on-target. First, regardless of MycER activation, cells treated with IACS-010759 for 24 hours showed a profound drop in mitochondrial processes, such as basal and maximal respiration, spare respiratory capacity and respiration-driven ATP production (Figure 2C). The same analysis revealed that MycER priming alone induced a statistically significant increase in both basal respiration and ATP production (Figure 2C), possibly associated with a moderate increase in the expression of OxPhos genes (supplemental Figure 2G). Second, ectopic expression of the *S. cerevisiae* Ndi1 protein, a IACS-010759-resistant ortholog of mammalian ETC complex I that can restore electron transport in complex I-deficient cells (16,29), conferred full resistance to IACS-010759 (supplemental Figure 3C).

The cytotoxic effects of IACS-010759 and other ETC disruptors were previously ascribed to a drop in cellular energy that could be compensated by increasing glycolytic rates, seemingly in line with the protective effect of glucose (28). However, energy depletion was unlikely to be the cause of cell death in our system, as neither OHT nor IACS-010759 impacted the ATP to ADP ratio, at any time before the onset of cell death (Figure 2D). This conservation of the energy balance following IACS-010759 treatment might be explained by adaptive cellular responses, such as increased glycolysis or suppression of energy-consuming processes (e. g. cell growth and proliferation). Mitochondrial respiration also sustains aspartate biosynthesis, which in turn is required for anabolic reactions, including protein and nucleotide biosynthesis (30,31). In line with this concept, supplementation with aspartate or nucleotide precursors partially bypassed the anti-proliferative effects of complex I inhibitors in other cell types (16,30–32). However, the same supplements were insufficient to prevent IACS-010759 induced arrest and cell death in FL^MycER^ and BaF^MycER^ cells (supplemental Figure 3D), implying that these effects involve additional complex I-dependent processes.

Cell death in FL^MycER^ cells treated with OHT and IACS-010759 presented clear features of apoptosis, such as external membrane exposure of phosphatidylserine (Annexin V staining: Figure 3A), chromatin condensation and nuclear fragmentation (supplemental Figure 4A), PARP cleavage (supplemental Figure 4B) and caspase activation – the latter suppressed by treatment with the caspase inhibitor Z-VAD-FMK (supplemental Figure 4C). Unexpectedly, however, Z-VAD-FMK failed to rescue IACS-010759-induced killing of OHT-primed cells (Figure 3B), implying that caspase activity in not an absolute requirement for apoptosis in this setting, as also observed in other contexts (33,34).

**Figure 4.**
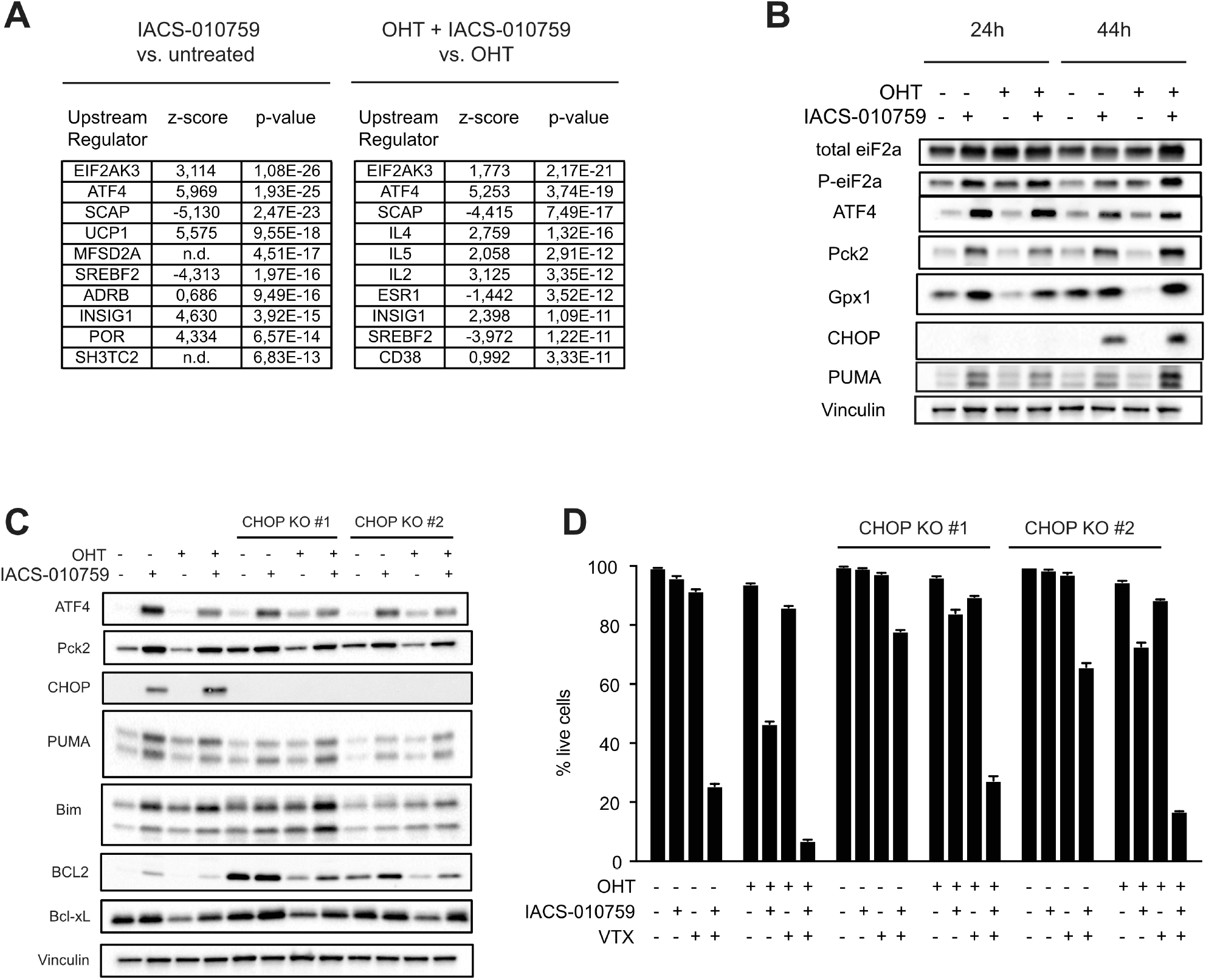
The Integrated Stress Response mediates IACS-010759-induced cell death. FL^MycER^ cells were pre-treated or not with OHT (100 nM, 48h), followed by IACS-010759 (135 nM, 24h). (**A**) Gene expression was profiled by RNA-seq, as defined in the methods section: the table shows the 10 most significantly enriched Upstream Regulators identified through analysis of IACS-010759-responsive genes, either with (right) or without OHT (left). n.d.: not determined. (**B**) Immunoblot analysis of ISR components in FL^MycER^ cells after treatment with IACS-010759 for 24 or 44h as indicated. Vinculin was used as loading control. (**C**) Immunoblot analysis of parental and CHOP-knockout FL^MycER^ cells after 44 hours of IACS-010759 treatment. (**D**) Percentage of live cells, confronting parental FL^MycER^ cells with two CHOP-knockout clones, treated with OHT, followed by IACS-010759 and/or venetoclax (VTX; 100 nM), as indicated. Error bars: SD (n=3).

The apoptotic response can be activated by either the extrinsic or the intrinsic pathway, the latter mediated by permeabilization of the mitochondrial outer membrane, determined by the equilibrium between pro- and anti-apoptotic members of the BCL2 protein family (35,36). In particular, the effectors Bax and Bak must disengage from anti-apoptotic BCL2-family proteins to form oligomeric pores, through which cytochrome c and other proteins are released from the mitochondrial intermembrane space into the cytosol (37). Indeed, we detected cytochrome c release from the mitochondria of OHT-primed FL^MycER^ cells undergoing IACS-010759-induced cell death (Figure 3C, supplemental Figure 4D). In order to further address the role of intrinsic apoptosis in our model, we derived Bak/Bak-null FL^MycER^ cell clones through CRISPR-Cas9 engineering: remarkably, these knockout cells acquired full resistance to IACS-010759-induced cell death (Figure 3D), while retaining the cytostatic response (Figure 3E). Hence, IACS-010759-mediated cytotoxicity was mediated by the intrinsic apoptotic pathway.

### BCL2 acts as a suppressor of IACS-010759-induced cell death

Myc is known to perturb the balance between pro- and anti-apoptotic BCL2-family members by promoting transcription of Bim (38,39) and Bax (40), while repressing the expression of BCL2 and Bcl-xL (41,42), altogether favoring cytochrome c release (43). In order to address whether the increased sensitivity to IACS-010759 following MycER activation could be ascribed to alterations in balance within the BCL2 family, we monitored protein levels in FL^MycER^ cells following OHT priming and subsequent treatment with IACS-010759 for 24 and 44 hours (both before the actual onset of cell death): indeed, MycER activation down-regulated BCL2 and Bcl-xL, over-riding a slight increase in these proteins promoted by IACS-010759 alone (Figure 3F). In contrast, neither Bim, nor Bax were significantly affected by MycER, while moderately induced by IACS-010759 (in particular at 44h: Figure 3F). Based on these results, we hypothesized that reduced expression of anti-apoptotic BCL2-family proteins upon MycER activation may cause the increased sensitivity to IACS-010759 in FL^MycER^ cells. In support of this concept, the BCL2-specific inhibitor venetoclax sensitized the cells to IACS-010759-induced cell death irrespective of prior MycER activation (Figure 3G), confirming the importance of endogenous BCL2 in conferring resistance to IACS-010759. Reciprocally, ectopic expression of BCL2 prevented IACS-010759-induced cell death in OHT-primed cells, a protective effect predictably overridden by venetoclax (Figure 3H).

Previous data indicated that BCL2 can sustain OxPhos activity in leukemic stem cells (44). Hence, in the above experiments, BCL2 might have acted by boosting mitochondrial respiration and/or protecting it from IACS-010759-mediated inhibition. In contrast with this scenario, however, neither over-expression, nor inhibition of BCL2 significantly impacted respiratory activity or its suppression by IACS-010759 in OHT-primed FL^MycER^ cells (supplemental Figure 4E, F). Finally, Bax/Bak-null FL^MycER^ cells remained resistant to IACS-010759-induced cytotoxicity even in the presence of venetoclax (supplemental Figure 4G), further emphasizing that BCL2 blocks IACS-010759-induced cell death by preventing activation of the intrinsic pathway.

### IACS-010759 treatment activates the Integrated Stress Response

In order to characterize possible signals mediating the cytotoxic action of IACS-010759, we profiled transcriptional changes after 24h of IACS-010759 treatment in either OHT-primed or non-primed FL^MycER^ cells. Independently from MycER activation, IACS-010759 elicited extensive transcriptional alterations, with over 1000 DEGs in either OHT-primed or non-primed cells and a close correlation between the two responses (supplemental Figure 3E). Upstream regulator analysis identified EIF2AK3 (also known as PERK) and ATF4, two key components of the Integrated Stress Response (ISR) (45), as controllers of the transcriptional response to IACS-010759 in both OHT-primed and non-primed cells (Figure 4A). Moreover, IACS-010759 treatment suppressed the expression of genes involved in sterol biosynthesis, which are controlled by the Sterol regulatory element-binding proteins (SREBPs) and the upstream SCAP-INSIG1 regulatory circuit (46). Finally, IACS-010759 led to the activation of genes associated with the expression of UCP1, which mediates dissipation of the proton gradient across the mitochondrial inner membrane, thus decoupling electron transport from ATP synthesis (47): this most likely indicates a common cellular response to impaired mitochondrial ATP production, regardless of its cause (i.e. suppression of membrane potential by UCP1, or disruption of the electron transport chain by IACS-010759; Figure 2C).

The ISR, the highest ranking IACS-010759-regulated program in our analysis, is an adaptive pathway engaged by diverse stress stimuli that selectively activate one of four different kinases (EIF2AK1-4), which converge to phosphorylate a specific residue (S51) in the translation factor eIF2a (45). While repressing general translation, phospho-eIF2a promotes the translation of a subset of transcripts, including the *ATF4* mRNA. ATF4 and several other ISR-induced transcriptional regulators, in turn, promote a pro-survival, stress-resistance program (45). Most relevant here, under conditions of severe, unresolved stress, the ISR can also induce pro-apoptotic factors such as the BH3-only BCL2-family proteins Bcl2l11/Bim, Bbc3/PUMA, Hrk and NOXA (48–52).

Based on the above, we monitored the status of the ISR pathway in FL^mycER^ cells: treatment with IACS-010759, but not OHT, promoted eIF2a phosphorylation and accumulation of ATF4 and several of its target-gene products, known to be involved in either cell survival (Gpx1, Pck2) or death (Ddit3/CHOP, PUMA, Bim) (Figure 4B and 3F) (45). Part of these were also significantly up-regulated at the mRNA level in our RNA-seq data (i.e. Pck2 and CHOP), while others were not (Gpx1, PUMA, Bim) (supplemental Figure 3F), possibly reflecting activation of the latter at the translational level, as described in other ISR-related contexts (53,54). Two other known effectors of the ISR, Hrk and NOXA (51), were not expressed in FL^mycER^ cells as judged by our RNA-seq data, precluding assessment of their possible roles in IACS-010759-induced cell death.

In order to assess the contribution of the ISR to the cytotoxic action of IACS-010759, we used CRISPR-Cas9 engineering to inactivate the gene encoding CHOP, the main mediator of the pro-apoptotic branch of the ISR (45). Previous work suggested that CHOP acts by inducing the BH3-only proteins PUMA and Bim (48–50), while suppressing BCL2 expression (55). However, similar to parental FL^MycER^ cells, CHOP-null cells showed IACS-010759-dependent activation of Bim and PUMA (Figure 4C). Instead, while still suppressed by MycER activation, BCL2 levels were altered in CHOP knock-out cells, with increased amounts in either basal or IACS-010759-treated conditions (Figure 4C). Consistent with the role of BCL2 in IACS-010759-induced cell death, CHOP KO cells showed increased resistance to the drug, which was overcome by co-treatment with venetoclax (Figure 4D).

### IACS-010759 and venetoclax synergize against MYC/BCL2 double-hit lymphoma

The preferential killing of Myc-overexpressing cells by IACS-010759 (Figure 2A) and its suppression by Bcl2 (Figure 3G, H) were reminiscent of our previous results with Tigecycline (11,20). We also reported that Tigecycline and venetoclax cooperated in killing *MYC/BCL2* double-hit lymphoma cells, and showed synergy against DHL in a pre-clinical setting (20). Hence, these observations prompted us to address the potential of combining IACS-010759 and venetoclax to treat DHL. Indeed, dosing either drug against the other *in vitro* on the human DHL cell lines SU-DHL-6 and DOHH-2 revealed a strong synergy in cell killing (Figure 5A, B), further substantiated with fixed concentrations of each drug in two other DHL lines (supplemental Figure 5A). Next, we addressed whether, as in FL5.12 cells (Figure 4A,B), IACS-010759 might also induce an ISR in DHL tumor cells. Indeed, IACS-010759 treatment of DHL cultures caused eIF2a phosphorylation and ATF4 accumulation (supplemental Figure 5B).

**Figure 5.**
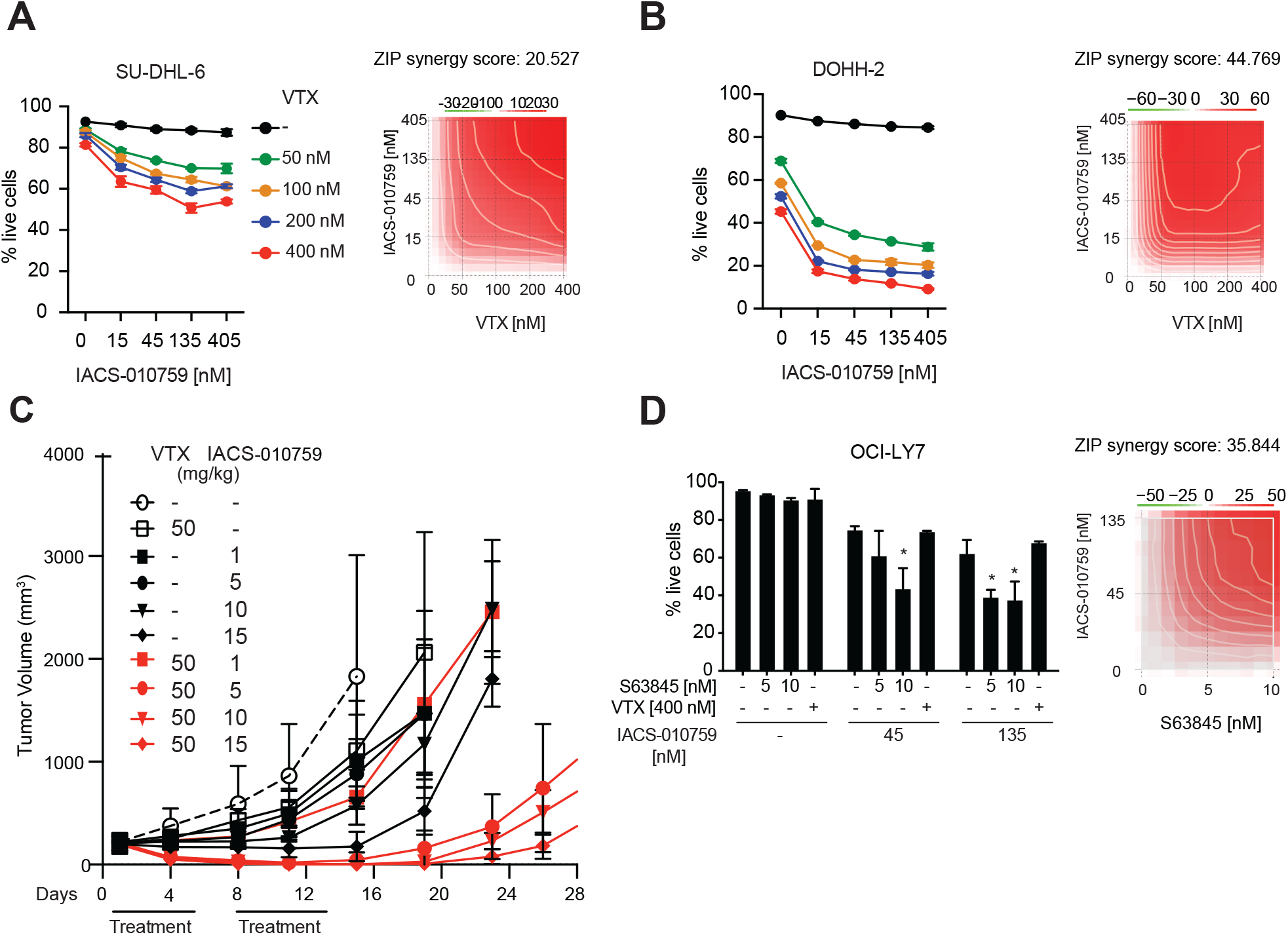
Combinatorial action of IACS-010759 and BH3-mimetic compounds against MYC-associated lymphomas. (**A**) Left: percentage of live SU-DHL-6 cells after 24 hours treatment with the indicated concentrations of IACS-010759 and venetoclax. Error bars: SD (n=3). Right: drug interaction landscape and synergy score for the two drugs calculated according to the ZIP model. Note that a positive ZIP score (>10) signifies a synergistic interaction. The landscape identifies the doses at which the drugs either synergize (red) or antagonize each-other (green) - the latter not observed here. (**B**) As in (A), for DOHH-2 cells. **(C)** Tumor progression in CD1 nude mice bearing subcutaneous SU-DHL-6 tumors treated by oral gavage with the indicated daily doses of IACS-010759 and/or venetoclax. Tumor volumes (mm^3^) were monitored at the indicated time-points. Error bars: SD; n = 5 animals per treatment group. p-values for the combinatorial treatments relative to untreated animals (white circles) at day 15 were the following: 0,0619 (50+1 group); 0,0098 (50+5); 0,0085 (50+10); 0,0185 (50+15). (**D**) OCI-LY7 lymphoma cells were treated for 24h with the indicated combinations of IACS-010759 with venetoclax or S63845. Left: percentage of live cells after treatment; error bars: SD (n=3). Right: drug interaction landscape and synergy score for IACS-010759 and S63845, calculated as in (A).

We then addressed the effects of IACS-010759 and venetoclax *in vivo* on subcutaneous SU-DHL-6 tumors in CD1 nude mice. As reported for tigecycline (20), we combined IACS-010759 (at doses between 1 and 15 mg/kg) with venetoclax (50mg/kg), delivered by daily oral gavage with 10 daily doses over two weeks: while either drug alone or the combination with the lowest dose of IACS-010759 (1 mg/kg) moderately slowed down tumor growth during the treatment period, combinations with higher doses of IACS-010759 showed increased efficacy and a marked delay in re-growth post-treatment (Figure 5C), most animals showing either partial or complete tumor regression up to one week after the end of treatment (supplemental Figure 5C: Expt. 1). These results were confirmed in a second experiment, with a similar response to the combination of venetoclax with IACS-010759 at 5 mg/kg, and a smaller, but significant response at 2.5 mg/kg (supplemental Figure 5C: Expt. 2). Hence, IACS-010759 and venetoclax provided strong anti-cancer effects against DHL in this pre-clinical setting.

The above results pointed to a wider therapeutic strategy against other high-grade lymphomas. In particular, in cases showing translocation of MYC alone, tumor progression and survival may be under the control of distinct anti-apoptotic BCL2-family proteins, against which specific pharmacological inhibitors are also available (21): the latter compounds – rather than venetoclax – may thus show synergy with IACS-010759 in these tumors. We tested this hypothesis in OCI-LY7, a BCL2-negative DLBCL cell line that is fully resistant to venetoclax (20), but expresses Mcl-1 (56). Indeed, while IACS-010759 alone showed substantial toxicity in those cells, this was significantly reinforced by co-treatment with the Mcl-1 specific inhibitor S63845, but not with venetoclax (Figure 5D). Finally, the Burkitt’s lymphoma cell line Ramos, which also expresses Mcl-1 but not BCL2, showed a similar sensitivity profile (supplemental Figure 5D). Thus, depending upon the cellular context, combination of IACS-010759 with distinct BH3-mimetic compounds may allow maximal anti-tumoral activity (Figure 6).

**Figure 6.**
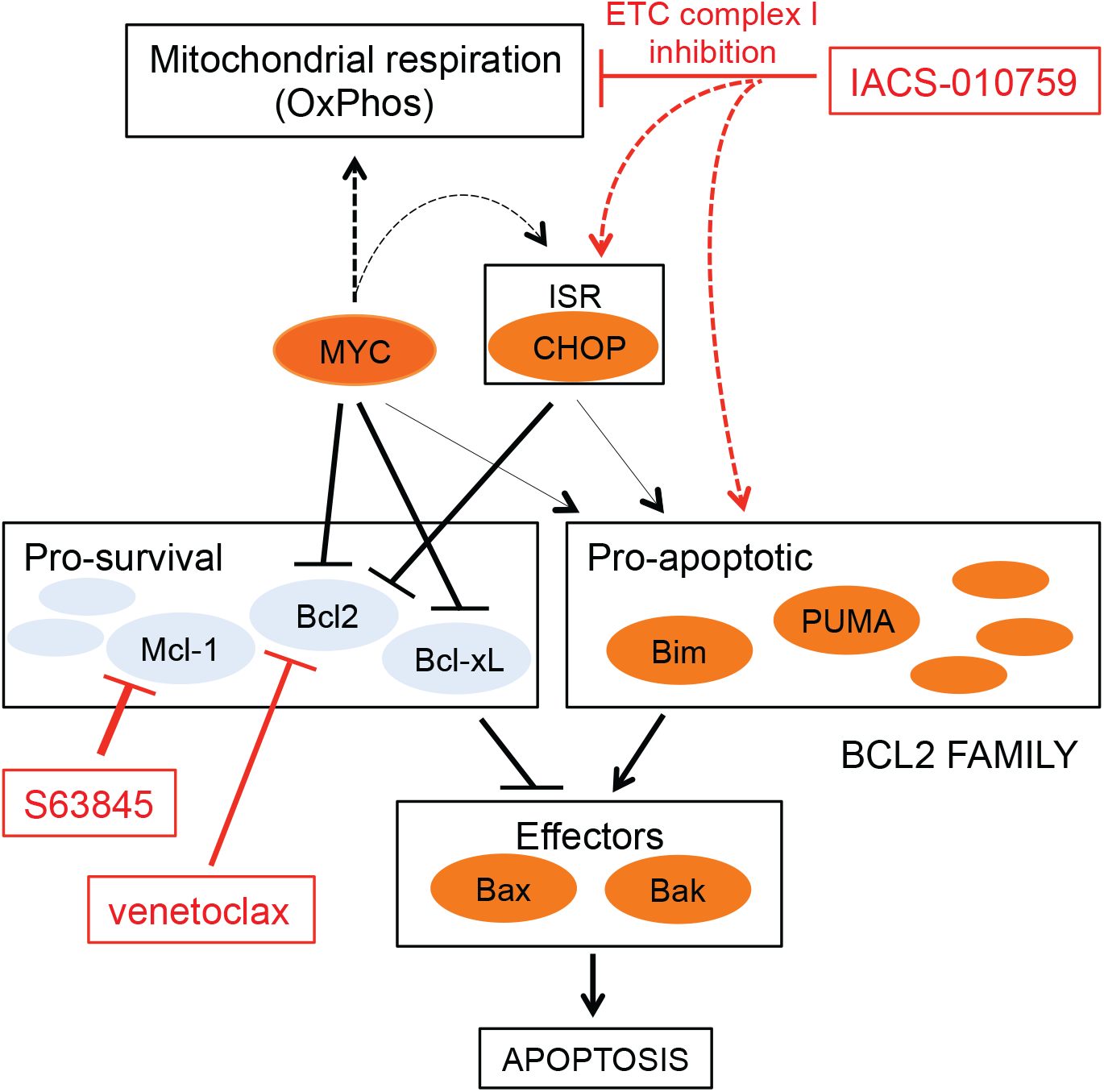
Combinatorial targeting of OxPhos and BCL2-family proteins in MYC-associated lymphoma. Schematic summary of the pharmaco-genetic interactions described in this work. Dashed arrows represent indirect effects; the connections between MYC, the integrated stress response (ISR) and the pro-apoptotic BCL2 arm (thin arrows) were described in other studies (see text), and may further reinforce the sensitivity of MYC-overexpressing cells to ETC inhibitors. Among a number of other pathways, MYC supports Mitochondrial respiration (Oxphos): inhibition of this process, and in particular of ETC complex I by IACS-010759 is synthetic-lethal with MYC, pointing to Oxphos as an important effector in MYC-induced tumorigenesis. Mechanistically, IACS-010759 induces apoptosis through activation of the ISR and in particular its pro-apoptotic effector CHOP, and may independently impact BCL2-family proteins. MYC sensitizes to apoptosis by modulating the expression of BCL2-family members (and, not shown here, activation of the ARF/p53 pathway). Hence, the ISR and MYC activity converge on the BCL2 family to lower the apoptotic threshold upon IACS-010759 treatment. This model provides a coherent rationale for the effects reported in this work, including (i.) MYC-induced sensitization of B-cells to killing by IACS-010759 and (ii.) the cooperative cytotoxic action of IACS-010759 and BH3-mimetic compounds, in particular venetoclax in MYC/BCL2 DHL cells, and S63845 in Mcl-1 expressing DLBCL and Burkitt’s lymphoma cells.

## Discussion

Our work points to Oxidative Phosphorylation (OxPhos) as one of the critical MYC-activated processes in DLBCL, and as a tractable therapeutic target in high-grade, MYC-associated forms of the disease. First, MYC- and OxPhos-related gene signatures were highly correlated in six distinct DLBCL patient cohorts, and were enriched in MYC-overexpressing mouse B-cells and lymphomas. Most noteworthy here, MYC may drive OxPhos genes not only at the transcriptional, but also at the translational level in B-cells (57). Second, ectopic MYC activity sensitized B-cells to IACS-010759, an inhibitor of ETC complex I (16). Third, IACS-010759 showed synergy with BH3-mimetic compounds that inhibit either BCL2 or Mcl-1, allowing context-dependent killing of lymphoma cell lines (Figure 6). Finally, IACS-010759 and the BCL2 inhibitor venetoclax effectively cooperated against *MYC/BCL2* double-hit lymphoma in a xenograft-based pre-clinical model.

It has long been known that MYC overexpression sensitizes diverse cell types to apoptosis in response to environmental or cell-autonomous stress conditions (27,43,58), in a manner that can be counteracted by BCL2 (58,59). In line with this concept, ectopic activation of a MycER chimera sensitized two mouse B-cell lines to killing by IACS-010759, an effect that was blocked by BCL2 and, reciprocally, exacerbated by venetoclax. Yet, unlike reported in AML cells (44,60), venetoclax did not suppress OxPhos activity in B-cells, indicating that it did not directly enhance IACS-010759 activity on ETC complex I. At the mechanistic level, and as described in other contexts (41,42), MycER activation suppressed expression of BCL2 and Bcl-xL. Further analysis confirmed that IACS-010759 induced cell death through the intrinsic apoptotic pathway, as evidenced by the requirement for the effectors Bax and Bak, and cytoplasmic release of cytochrome c. Altogether, as depicted in Figure 6, MYC promoted OxPhos activity and concomitantly lowered the anti-apoptotic safeguard provided by BCL2/Bcl-xL in B-cells (61), thus sensitizing the cells to IACS-010759.

To identify possible effectors of IACS-010759, we profiled gene expression in B-cells: this singled out the Integrated Stress Response (ISR) as a IACS-010759-induced pathway, independently from MycER. The ISR is a composite signaling pathway that mediates protective adaptation to multiple stresses, but concurrently promotes cell death when homeostasis cannot be restored (45): activation of this pathway in response to OxPhos inhibitors was also reported in AML, multiple myeloma and glioblastoma cells, in which it appeared to relay mainly a pro-death signal (60,62,63). In line with these observations, elimination of CHOP, which controls the pro-apoptotic branch of the ISR (45), conferred resistance to IACS-010759 in our experiments, thus pointing to the ISR as a common effector of OxPhos inhibitors in diverse tumor types. Finally, we shall note here that MYC-induced oncogenic stress can also activate the ISR (64): while not observed in our experimental setting, this may further contribute to the sensitivity of MYC-driven tumors to ETC inhibitors (Figure 6).

While IACS-010759 directly inhibits ETC complex I, thus suppressing OxPhos activity (16), tigecycline achieves the same effect indirectly via inhibition of mitochondrial translation (11,13), providing a common denominator for their pro-apoptotic effects on MYC-overexpressing cells, as well as on DHL cells when combined with venetoclax (20). Thus, targeting OxPhos (whether with IACS-010759, tigecycline, etc…) along with select BCL2-family members (21) may be an effective means to achieve synergy against high-grade B-cell lymphomas (Figure 6). In support of this concept, the BCL2 inhibitor venetoclax potentiated the cytotoxic activity of IACS-010759 in *MYC/BCL2* DHL cells, while the Mcl-1 inhibitor S63845 did so in MYC-translocated, BCL2-negative lymphoma cell lines. Altogether, our work supports a wider therapeutic concept, in which the status of MYC and BCL2 family members in individual patients may guide the decision to combine OxPhos inhibitors and select BH3-mimetics against high-grade DLBCL, and possibly other refractory malignancies.

## Acknowledgments

We thank Simona Ronzoni for support with flow cytometry, Salvatore Bianchi, Luca Rotta and Thelma Capra for support with the Illumina HiSeq platform, Alberto Gobbi and Manuela Capillo for their help with the management of mouse colonies and Ottavio Croci for guidance on data analysis, Andreas Strasser, Marco Herald and Grant Dewson for advice on CRISPR-Cas9 targeting of mouse *Bax* and *Bak*, Andrea Viale for the initial suggestion to use IACS-010759, Marzia Fumagalli and Claudia Iavarone for help with material transfer agreements, Stefano Campaner, Saverio Minucci, Riccardo Cazzoli, Gioacchino Natoli and members of the Amati lab for insightful comments and discussions. We are grateful to Tushar Dave and Anupama Reddy for providing us access to the EGAS00001002606 dataset [ref. (3)], under an agreement with the Duke University (Durham NC, USA), and to Stacy Hung, Kal Mann, Theolina Dimitrow and Christian Steidl for the EGAS00001002657 dataset [ref. (22)], under an agreement with BC Cancer (Vancouver BC, Canada); the results and conclusions reported in this work do not necessarily reflect the opinions or views of BC Cancer. The dataset phs001444 [ref. (5)] was accessed with a Data Use Certification Agreement from the NIH database for Genotypes and Phenotypes (dbGaP).

## Author contributions

GD and MR organized and performed most of the experiments, with technical assistance from PN, AV and MD. MF performed the computational analyses. FP contributed part of the *in vitro* experiments on cellular drug responses. LC and AB provided support and assistance with the SeaHorse experiments, SR with microscopy, and CPV, JRM and GFD with the provision of IACS-010759. GD and BA designed and coordinated the project, and wrote the manuscript.

## References

1. Liu Y, Barta SK. Diffuse large B-cell lymphoma: 2019 update on diagnosis, risk stratification, and treatment. Am J Hematol 2019;94:604–16

2. Monti S, Savage KJ, Kutok JL, Feuerhake F, Kurtin P, Mihm M, et al. Molecular profiling of diffuse large B-cell lymphoma identifies robust subtypes including one characterized by host inflammatory response. Blood 2005;105:1851–61

3. Reddy A, Zhang J, Davis NS, Moffitt AB, Love CL, Waldrop A, et al. Genetic and Functional Drivers of Diffuse Large B Cell Lymphoma. Cell 2017;171:481–94 e15

4. Chapuy B, Stewart C, Dunford AJ, Kim J, Kamburov A, Redd RA, et al. Molecular subtypes of diffuse large B cell lymphoma are associated with distinct pathogenic mechanisms and outcomes. Nat Med 2018;24:679–90

5. Schmitz R, Wright GW, Huang DW, Johnson CA, Phelan JD, Wang JQ, et al. Genetics and Pathogenesis of Diffuse Large B-Cell Lymphoma. N Engl J Med 2018;378:1396–407

6. Wright GW, Huang DW, Phelan JD, Coulibaly ZA, Roulland S, Young RM, et al. A Probabilistic Classification Tool for Genetic Subtypes of Diffuse Large B Cell Lymphoma with Therapeutic Implications. Cancer Cell 2020;37:551–68 e14

7. Bisso A, Sabò A, Amati B. MYC in Germinal Center-derived lymphomas: Mechanisms and therapeutic opportunities. Immunol Rev 2019;288:178–97

8. Chiche J, Reverso-Meinietti J, Mouchotte A, Rubio-Patino C, Mhaidly R, Villa E, et al. GAPDH Expression Predicts the Response to R-CHOP, the Tumor Metabolic Status, and the Response of DLBCL Patients to Metabolic Inhibitors. Cell Metab 2019;29:1243–57 e10

9. Carey CD, Gusenleitner D, Chapuy B, Kovach AE, Kluk MJ, Sun HH, et al. Molecular classification of MYC-driven B-cell lymphomas by targeted gene expression profiling of fixed biopsy specimens. J Mol Diagn 2015;17:19–30

10. Morrish F, Hockenbery D. MYC and mitochondrial biogenesis. Cold Spring Harb Perspect Med 2014;4

11. D’Andrea A, Gritti I, Nicoli P, Giorgio M, Doni M, Conti A, et al. The mitochondrial translation machinery as a therapeutic target in Myc-driven lymphomas. Oncotarget 2016;7:72415–30

12. Oran AR, Adams CM, Zhang XY, Gennaro VJ, Pfeiffer HK, Mellert HS, et al. Multi-focal control of mitochondrial gene expression by oncogenic MYC provides potential therapeutic targets in cancer. Oncotarget 2016;7:72395–414

13. Skrtic M, Sriskanthadevan S, Jhas B, Gebbia M, Wang X, Wang Z, et al. Inhibition of mitochondrial translation as a therapeutic strategy for human acute myeloid leukemia. Cancer Cell 2011;20:674–88

14. Dong Z, Abbas MN, Kausar S, Yang J, Li L, Tan L, et al. Biological Functions and Molecular Mechanisms of Antibiotic Tigecycline in the Treatment of Cancers. Int J Mol Sci 2019;20

15. Norberg E, Lako A, Chen PH, Stanley IA, Zhou F, Ficarro SB, et al. Differential contribution of the mitochondrial translation pathway to the survival of diffuse large B-cell lymphoma subsets. Cell Death Differ 2017;24:251–62

16. Molina JR, Sun YT, Protopopova M, Gera S, Bandi M, Bristow C, et al. An inhibitor of oxidative phosphorylation exploits cancer vulnerability. Nat Med 2018;24:1036–46

17. Vangapandu HV, Alston B, Morse J, Ayres ML, Wierda WG, Keating MJ, et al. Biological and metabolic effects of IACS-010759, an OxPhos inhibitor, on chronic lymphocytic leukemia cells. Oncotarget 2018;9:24980–91

18. Zhang L, Yao Y, Zhang S, Liu Y, Guo H, Ahmed M, et al. Metabolic reprogramming toward oxidative phosphorylation identifies a therapeutic target for mantle cell lymphoma. Sci Transl Med 2019;11

19. Deribe YL, Sun Y, Terranova C, Khan F, Martinez-Ledesma J, Gay J, et al. Mutations in the SWI/SNF complex induce a targetable dependence on oxidative phosphorylation in lung cancer. Nat Med 2018;24:1047–57

20. Ravà M, D’Andrea A, Nicoli P, Gritti I, Donati G, Doni M, et al. Therapeutic synergy between tigecycline and venetoclax in a preclinical model of MYC/BCL2 double-hit B cell lymphoma. Sci Transl Med 2018;10

21. Adams CM, Clark-Garvey S, Porcu P, Eischen CM. Targeting the Bcl-2 Family in B Cell Lymphoma. Front Oncol 2018;8:636

22. Ennishi D, Jiang A, Boyle M, Collinge B, Grande BM, Ben-Neriah S, et al. Double-Hit Gene Expression Signature Defines a Distinct Subgroup of Germinal Center B-Cell-Like Diffuse Large B-Cell Lymphoma. J Clin Oncol 2019;37:190–201

23. Lenz G, Wright GW, Emre NC, Kohlhammer H, Dave SS, Davis RE, et al. Molecular subtypes of diffuse large B-cell lymphoma arise by distinct genetic pathways. Proc Natl Acad Sci U S A 2008;105:13520–5

24. Sha C, Barrans S, Cucco F, Bentley MA, Care MA, Cummin T, et al. Molecular High-Grade B-Cell Lymphoma: Defining a Poor-Risk Group That Requires Different Approaches to Therapy. J Clin Oncol 2019;37:202–12

25. Sabò A, Kress TR, Pelizzola M, de Pretis S, Gorski MM, Tesi A, et al. Selective transcriptional regulation by Myc in cellular growth control and lymphomagenesis. Nature 2014;511:488–92

26. Schleich K, Kase J, Dörr JR, Trescher S, Bhattacharya A, Yu Y, et al. H3K9me3-mediated epigenetic regulation of senescence in mice predicts outcome of lymphoma patients. Nature Communications 2020;In press

27. Evan GI, Wyllie AH, Gilbert CS, Littlewood TD, Land H, Brooks M, et al. Induction of apoptosis in fibroblasts by c-myc protein. Cell 1992;69:119–28

28. Naguib A, Mathew G, Reczek CR, Watrud K, Ambrico A, Herzka T, et al. Mitochondrial Complex I Inhibitors Expose a Vulnerability for Selective Killing of Pten-Null Cells. Cell Rep 2018;23:58–67

29. Seo BB, Kitajima-Ihara T, Chan EKL, Scheffler IE, Matsuno-Yagi A, Yagi T. Molecular remedy of complex I defects: Rotenone-insensitive internal NADH-quinone oxidoreductase of Saccharomyces cerevisiae mitochondria restores the NADH oxidase activity of complex I-deficient mammalian cells. P Natl Acad Sci USA 1998;95:9167–71

30. Birsoy K, Wang T, Chen WW, Freinkman E, Abu-Remaileh M, Sabatini DM. An Essential Role of the Mitochondrial Electron Transport Chain in Cell Proliferation Is to Enable Aspartate Synthesis. Cell 2015;162:540–51

31. Sullivan LB, Gui DY, Hosios AM, Bush LN, Freinkman E, Vander Heiden MG. Supporting Aspartate Biosynthesis Is an Essential Function of Respiration in Proliferating Cells. Cell 2015;162:552–63

32. Sullivan LB, Luengo A, Danai LV, Bush LN, Diehl FF, Hosios AM, et al. Aspartate is an endogenous metabolic limitation for tumour growth. Nat Cell Biol 2018;20:782–8

33. Kroemer G, Martin SJ. Caspase-independent cell death. Nat Med 2005;11:725–30

34. Tait SW, Green DR. Mitochondrial regulation of cell death. Cold Spring Harb Perspect Biol 2013;5

35. Czabotar PE, Lessene G, Strasser A, Adams JM. Control of apoptosis by the BCL-2 protein family: implications for physiology and therapy. Nat Rev Mol Cell Biol 2014;15:49–63

36. Kalkavan H, Green DR. MOMP, cell suicide as a BCL-2 family business. Cell Death Differ 2018;25:46–55

37. Luna-Vargas MPA, Chipuk JE. Physiological and Pharmacological Control of BAK, BAX, and Beyond. Trends Cell Biol 2016;26:906–17

38. Campone M, Noel B, Couriaud C, Grau M, Guillemin Y, Gautier F, et al. c-Myc dependent expression of pro-apoptotic Bim renders HER2-overexpressing breast cancer cells dependent on anti-apoptotic Mcl-1. Mol Cancer 2011;10:110

39. Muthalagu N, Junttila MR, Wiese KE, Wolf E, Morton J, Bauer B, et al. BIM is the primary mediator of MYC-induced apoptosis in multiple solid tissues. Cell Rep 2014;8:1347–53

40. Mitchell KO, Ricci MS, Miyashita T, Dicker DT, Jin Z, Reed JC, et al. Bax is a transcriptional target and mediator of c-myc-induced apoptosis. Cancer Res 2000;60:6318–25

41. Eischen CM, Woo D, Roussel MF, Cleveland JL. Apoptosis triggered by Myc-induced suppression of Bcl-X(L) or Bcl-2 is bypassed during lymphomagenesis. Mol Cell Biol 2001;21:5063–70

42. Maclean KH, Keller UB, Rodriguez-Galindo C, Nilsson JA, Cleveland JL. c-Myc augments gamma irradiation-induced apoptosis by suppressing Bcl-XL. Mol Cell Biol 2003;23:7256–70

43. Juin P, Hueber AO, Littlewood T, Evan G. c-Myc-induced sensitization to apoptosis is mediated through cytochrome c release. Genes Dev 1999;13:1367–81

44. Lagadinou ED, Sach A, Callahan K, Rossi RM, Neering SJ, Minhajuddin M, et al. BCL-2 inhibition targets oxidative phosphorylation and selectively eradicates quiescent human leukemia stem cells. Cell Stem Cell 2013;12:329–41

45. Pakos-Zebrucka K, Koryga I, Mnich K, Ljujic M, Samali A, Gorman AM. The integrated stress response. EMBO Rep 2016;17:1374–95

46. McPherson R, Gauthier A. Molecular regulation of SREBP function: the Insig-SCAP connection and isoform-specific modulation of lipid synthesis. Biochem Cell Biol 2004;82:201–11

47. Klaus S, Casteilla L, Bouillaud F, Ricquier D. The uncoupling protein UCP: a membraneous mitochondrial ion carrier exclusively expressed in brown adipose tissue. Int J Biochem 1991;23:791–801

48. Puthalakath H, O’Reilly LA, Gunn P, Lee L, Kelly PN, Huntington ND, et al. ER stress triggers apoptosis by activating BH3-only protein Bim. Cell 2007;129:1337–49

49. Galehdar Z, Swan P, Fuerth B, Callaghan SM, Park DS, Cregan SP. Neuronal apoptosis induced by endoplasmic reticulum stress is regulated by ATF4-CHOP-mediated induction of the Bcl-2 homology 3-only member PUMA. J Neurosci 2010;30:16938–48

50. Ghosh AP, Klocke BJ, Ballestas ME, Roth KA. CHOP potentially co-operates with FOXO3a in neuronal cells to regulate PUMA and BIM expression in response to ER stress. PLoS One 2012;7:e39586

51. Pike LR, Phadwal K, Simon AK, Harris AL. ATF4 orchestrates a program of BH3-only protein expression in severe hypoxia. Mol Biol Rep 2012;39:10811–22

52. Wali JA, Rondas D, McKenzie MD, Zhao Y, Elkerbout L, Fynch S, et al. The proapoptotic BH3-only proteins Bim and Puma are downstream of endoplasmic reticulum and mitochondrial oxidative stress in pancreatic islets in response to glucotoxicity. Cell Death Dis 2014;5:e1124

53. Albert AE, Adua SJ, Cai WL, Arnal-Estape A, Cline GW, Liu Z, et al. Adaptive Protein Translation by the Integrated Stress Response Maintains the Proliferative and Migratory Capacity of Lung Adenocarcinoma Cells. Mol Cancer Res 2019;17:2343–55

54. Sidrauski C, McGeachy AM, Ingolia NT, Walter P. The small molecule ISRIB reverses the effects of eIF2alpha phosphorylation on translation and stress granule assembly. Elife 2015;4

55. McCullough KD, Martindale JL, Klotz LO, Aw TY, Holbrook NJ. Gadd153 sensitizes cells to endoplasmic reticulum stress by down-regulating Bcl2 and perturbing the cellular redox state. Mol Cell Biol 2001;21:1249–59

56. Klanova M, Andera L, Brazina J, Svadlenka J, Benesova S, Soukup J, et al. Targeting of BCL2 Family Proteins with ABT-199 and Homoharringtonine Reveals BCL2- and MCL1-Dependent Subgroups of Diffuse Large B-Cell Lymphoma. Clin Cancer Res 2016;22:1138–49

57. Singh K, Lin J, Zhong Y, Burcul A, Mohan P, Jiang M, et al. c-MYC regulates mRNA translation efficiency and start-site selection in lymphoma. J Exp Med 2019;216:1509–24

58. Bissonnette RP, Echeverri F, Mahboubi A, Green DR. Apoptotic cell death induced by c-myc is inhibited by bcl-2. Nature 1992;359:552–4

59. Fanidi A, Harrington EA, Evan GI. Cooperative interaction between c-myc and bcl-2 proto-oncogenes. Nature 1992;359:554–6

60. Sharon D, Cathelin S, Mirali S, Di Trani JM, Yanofsky DJ, Keon KA, et al. Inhibition of mitochondrial translation overcomes venetoclax resistance in AML through activation of the integrated stress response. Sci Transl Med 2019;11

61. Marsden VS, Strasser A. Control of apoptosis in the immune system: Bcl-2, BH3-only proteins and more. Annu Rev Immunol 2003;21:71–105

62. Bajpai R, Sharma A, Achreja A, Edgar CL, Wei C, Siddiqa AA, et al. Electron transport chain activity is a predictor and target for venetoclax sensitivity in multiple myeloma. Nat Commun 2020;11:1228

63. Shi Y, Lim SK, Liang Q, Iyer SV, Wang HY, Wang Z, et al. Gboxin is an oxidative phosphorylation inhibitor that targets glioblastoma. Nature 2019;567:341–6

64. Tameire F, Verginadis, II, Leli NM, Polte C, Conn CS, Ojha R, et al. ATF4 couples MYC-dependent translational activity to bioenergetic demands during tumour progression. Nat Cell Biol 2019;21:889–99

