## Supplemental Methods, Tables and Figures for "Targeting mitochondrial respiration and the BCL2 family in MYC-associated B-cell lymphoma"

For:

### Supplemental Materials and Methods

*Cell lines*

The murine lymphoid precursor cell lines Ba/F3 and FL5.12 (RRID: CVCL_0161 and CVCL _0262) (1,2) and their derivatives were grown in regular RPMI medium (Euroclone, Pero, Italy), which includes 2mM glutamine and 11 mM glucose, supplemented with 10% fetal bovine serum (FBS) and murine interleukin 3 (PeproTech, Rocky Hill, NJ, USA) at a final concentration of 1 and 2 ng/mL, respectively. The human lymphoma cell lines DOHH-2, SU-DHL-6, SU-DHL-4, Karpas 422, OCI-LY7 and Ramos (RRID: CVCL_1179, CVCL_2206, CVCL_0539, CVCL_1325, CVCL_1881 and CVCL_0597) were maintained in RPMI medium (including 11 mM glucose) supplemented with 10% FBS. Prior to experiments involving IACS-010759, all cells were passaged in glucose-free RPMI-1640 medium (Thermo Fisher Scientific, Waltham, MA, USA), which includes 2mM glutamine, supplemented with 10% FBS and 2.75 mM glucose. All cells were incubated at 37° C in a humidified air atmosphere supplemented with 5% CO2. OCI-LY7 and Ramos were selected to address differential sensitivity to the Mcl-1 inhibitor S63845 since they express Mcl-1 but not BCL2, and are thus resistant to venetoclax (3-6).

*Retroviral vectors*

Stable expression of exogenous proteins in Ba/F3 and FL5.12 cells was achieved by infection with retroviral vectors. In particular, MycER$^{T2}$ (here MycER) (7) was expressed with the pBabe-bleo vector (8), human BCL2 with a pMSCV Puromycin vector (9) (a gift from Joseph Opferman), and Ndi1 with PMXS-Blasticidin (Addgene, Watertwon, CA, USA; plasmid # 72876, a gift from David Sabatini). Transduced cells were selected for one week with the appropriate antibiotic at the following concentrations: zeocin 2 µg/mL, puromycin 1.5 µg/mL, blasticidin 5µg/mL.

*Reagents and kits*

L-aspartic acid (Merck, Darmstadt, Germany) was added to the growth medium and the pH adjusted to 7.5 with 1 M NaOH before FBS supplementation; hypoxanthine, adenine and uridine (all from Merck Life Science) were solubilized and added directly to the growth medium. IACS-010759 (10), venetoclax/ABT-199 (Carbosynth, Newbury, UK) (11), S63845 (Medchemexpress LCC, Monmouth Junction, NJ, USA) (12), Z-VAD-FMK (Santa Cruz Biotechnologies, Santa Cruz, CA, USA) were dissolved in DMSO and added directly to the culture medium; 4-hydroxytamoxifen (OHT; Merck) was dissolved in ethanol and added directly to the culture medium.

An ADP/ATP Ratio Assay kit (Merck) was used to quantify cellular ADP and ATP, while Caspase-Glo 3/7 Assay (Promega, Madison, WI, USA) was used to evaluate caspase activity, both according to the manufacturer's instructions.

*Immunoblot analysis*

After collection, cells were divided in 2 aliquots: one was extracted as previously described (13) to be used for quantification by Bradford protein assay (Bio-Rad Laboratories, Hercules, CA, USA), while the other was directly lysed in Laemmli buffer and denatured at 95 °C for 5 minutes for subsequent SDS-PAGE. After clearance by centrifugation, 10 µg of proteins from each sample were loaded on an acrylamide gel for SDS-PAGE, followed by transfer on a nitrocellulose membrane and immunoblot with the antibodies listed in Supplemental Table 2.

*Fluorescence microscopy*

Non-adherent cells were subjected to cytospin, fixed in 4% paraformaldehyde (PFA), and permeabilized with 0.1% Triton X-100 for subsequent staining as described previously (14). Single optical sections were acquired with an SP8 confocal microscope (Leica Microsystems, Wetzlar, Germany) equipped with a 63x/1.4 oil immersion objective lens. For the analysis of apoptotic figures, cells were stained with 4′,6-diamidino-2-phenylindole (DAPI) and tetramethylrhodamine (TRITC) -conjugated Agglutinin (both from Thermo Fisher Scientific, Waltham, MA, USA). For cytochrome c localization, cells were stained overnight at 4°C with an antibody against cytochrome c (clone 7H8.2C12, Thermo Fisher) and counter-stained with DAPI. Secondary staining was performed with an Alexa Fluor 488 -conjugated anti-mouse antibody (Jackson ImmunoResearch, West Grove, PA, USA). The total and cytochrome c-positive cellular areas were calculated from >20 fields for each sample with an in-house developed macro for ImageJ (RRID: SCR_003070). Briefly, the cytochrome c signal from each cell in the field of view was segmented using an automatic threshold (Otsu algorithm) and the area of the signal calculated on the binary images. The cytochrome c area was then normalized by the total cell area, which was identified on the bright-field transmission image.

*Flow Cytometry*

Flow cytometry was conducted on a MACSQuant Analyzer (Milteniy Biotec, Bergisch Gladbach, Germany) and data analyzed with FlowJo software (version 10.6.1; BD Biosciences, Franklin Lakes, NK, USA; RRID: SCR_008520). For cell number and viability counts, cells were resuspended in ice-cold PBS in the presence of 1 μg/mL Propidium Iodide (P.I.). Cell viability is given as percentage of live cells on total cells counted, while the Proliferation index is defined as the ratio of viable cells in the sample to viable cells in the corresponding IACS-010759-untreated control.

For evaluation of apoptotic cell death, cells were stained with APC-conjugated Annexin V (Thermo Fisher Scientific) and P.I. as previously described (15). The cytoplasmic release of cytochrome c was detected as described (16) with few modifications: briefly, cells were permeabilized with 0.002% digitonin for 20” under vortexing before fixation in 4% PFA and permeabilization with 0.1% Triton X-100, for subsequent immunostaining with anti-cytochrome c (clone 6H2.B4, BD Biosciences) overnight at 4°C. Secondary staining was performed with an Alexa Fluor 647 -conjugated anti-mouse antibody (Jackson ImmunoResearch).

*Cell cycle kinetics*

For the analysis of cell cycle progression, cells were incubated for 20 minutes in the presence of 33 μM bromodeoxyuridine (BrdU, Merck), washed, resuspended and incubated in fresh medium: at the indicated time-points (supplemental Figure 2A), aliquots were collected for ethanol-fixation followed by FITC-conjugated anti-BrdU (BD Biosciences) and P.I. staining. Stained samples were analyzed by flow cytometry on a MACSQuant Analyzer.

*RNA extraction and quantitative PCR analysis*

Total cellular RNA was extracted using the Quick-RNA Miniprep kit (Zymo Research, Irvine, CA, USA) and reverse transcribed with the iSCRIPT cDNA Synthesis Kit (Bio-Rad Laboratories). 10 ng of cDNA were used as template in each real-time quantitative PCR (qPCR) reaction, performed with fast SyberGreen Master Mix (Thermo Fisher Scientific) on a CFX96 Touch Real-Time PCR Detection System (Bio-Rad Laboratories). Primer sequences are provided in Supplemental Table 3.

*RNA library preparation and Next Generation Sequencing*

Libraries for RNA-Seq were prepared for each sample from 0.5 μg of total RNA treated with the Ribozero rRNA removal kit (Illumina, San Diego, CA, USA) and ethanol-precipitated. RNA quality and removal of rRNA were checked with the Agilent 2100 Bioanalyzer (Agilent Technologies, Santa Clara, CA, USA). Libraries for RNA-Seq were then prepared with the TruSeq RNA Sample Prep Kits v2 (Illumina) following the manufacturer's instructions (skipping the mRNA purification step with poly-T oligo-attached magnetic beads). RNA-Seq libraries were then run on the Agilent 2100 Bioanalyzer for quantification and quality control, and then sequenced on HiSeq2000 (Illumina). RNA-Seq reads were filtered using the fastq quality trimmer and fastq masker tools of the FASTX-Toolkit suite (http://hannonlab.cshl.edu/fastx_toolkit/; RRID: SCR_005534). Their quality was evaluated and confirmed using the FastQC application (http://www.bioinformatics.babraham.ac.uk/projects/fastqc/; RRID: SCR_014583). Pipelines for primary analysis (filtering and alignment to the reference genome of the raw reads) and secondary analysis (expression quantification, differential gene expression, and peak calling) have been integrated in the HTS-flow system (17). Bioinformatic and statistical analysis were performed using R with Bioconductor packages (RRID: SCR_006442) (18). Differentially expressed genes (DEGs) were identified using the Bioconductor Deseq2 package (RRID: SCR_015687) (19) as genes whose q-value is lower than 0.05. Gene set enrichment analysis (GSEA) was performed using the Desktop tool of the Broad Institute (https://www.gsea-msigdb.org/gsea/index.jsp; RRID: SCR_003199) (20) by querying DEGs for the enrichment of Hallmark Gene Sets from the Hallmark MSigDB collection (21) and CCC DLBCL signatures (22). Upstream regulator analysis of DEGs was performed with the Ingenuity Pathway Analysis software package (QIAGEN, Venlo, Netherlands; RRID: SCR_008653). The RNA-Seq data described in this work are accessible through NCBI's Gene Expression Omnibus (GEO; RRID: SCR_005012) (23) with series accession numbers GSE149073 and GSE51011.

*CRISPR-Cas9 constructs and knockout generation*

Gene knockout was obtained by targeting two sites for each gene, selected using the CRISPR Targets Track function from UCSC Genome Browser (https://genome-euro.ucsc.edu/; RRID: SCR_005780): one was selected close to, and the other ca. 100 bases downstream of the start codon. The genomic sequences targeted by the corresponding sgRNAs are provided in Supplemental Table 4. Complementary DNA oligonucleotides encompassing the sequence of each sgRNA were annealed and ligated into the PX458 plasmid (Addgene; plasmid # 48138, a gift from Feng Zhang) digested with BbsI, as described (24). For each construct, 1 µg of DNA was electroporated in $4x10^5$ FL5.12 cells using the Neon Transfection System (Thermo Fisher Scientific). After two days, GFP positive cells were sorted with FACSMelody (BD Biosciences) and single clones isolated by limiting dilution. Following in vitro expansion, each clone was tested for recombination by extracting genomic DNA with the QuickExtract DNA Extraction Solution (Illumina) and performing PCR amplification with

GoTaq polymerase (Promega) with specific primers for each genomic locus (Supplemental Table 5). Ablation of the targeted protein in the selected clones was confirmed by immunoblot analysis.

*Quantification and statistical analysis*

Drug interaction landscapes and delta scores for synergy were based on the ZIP model in SynergyFinder (25). All other statistical analyses were performed with GraphPad PRISM 8 (RRID: SCR_002798); multiple comparisons between groups from cell culture experiments were performed by one-way ANOVA with Dunnet's test. The numbers of independent biological replicates in cell culture experiments are indicated in the corresponding figures. Statistical evaluation of tumor growth (Figure 5C) was performed using multiple t tests for each time point and the Holm-Sidak method.

### Supplemental References


1. Palacios R, Steinmetz M. Il-3-dependent mouse clones that express B-220 surface antigen, contain Ig genes in germ-line configuration, and generate B lymphocytes in vivo. Cell **1985**;41:727-34
2. Mckearn JP, Mccubrey J, Fagg B. Enrichment of Hematopoietic Precursor Cells and Cloning of Multipotential Lymphocyte-B Precursors. P Natl Acad Sci USA **1985**;82:7414-8
3. Ravà M, D'Andrea A, Nicoli P, Gritti I, Donati G, Doni M, *et al.* Therapeutic synergy between tigecycline and venetoclax in a preclinical model of MYC/BCL2 double-hit B cell lymphoma. Sci Transl Med **2018**;10
4. Johnson-Farley N, Veliz J, Bhagavathi S, Bertino JR. ABT-199, a BH3 mimetic that specifically targets Bcl-2, enhances the antitumor activity of chemotherapy, bortezomib and JQ1 in "double hit" lymphoma cells. Leuk Lymphoma **2015**;56:2146-52
5. Stolz C, Hess G, Hahnel PS, Grabellus F, Hoffarth S, Schmid KW*, et al.* Targeting Bcl-2 family proteins modulates the sensitivity of B-cell lymphoma to rituximab-induced apoptosis. Blood **2008**;112:3312-21
6. Klanova M, Andera L, Brazina J, Svadlenka J, Benesova S, Soukup J*, et al.* Targeting of BCL2 Family Proteins with ABT-199 and Homoharringtonine Reveals BCL2- and MCL1-Dependent Subgroups of Diffuse Large B-Cell Lymphoma. Clin Cancer Res **2016**;22:1138-49
7. Littlewood TD, Hancock DC, Danielian PS, Parker MG, Evan GI. A modified oestrogen receptor ligand-binding domain as an improved switch for the regulation of heterologous proteins. Nucleic Acids Res **1995**;23:1686-90
8. Morgenstern JP, Land H. Advanced mammalian gene transfer: high titre retroviral vectors with multiple drug selection markers and a complementary helper-free packaging cell line. Nucleic Acids Res **1990**;18:3587-96
9. Koss B, Ryan J, Budhraja A, Szarama K, Yang X, Bathina M*, et al.* Defining specificity and on-target activity of BH3-mimetics using engineered B-ALL cell lines. Oncotarget **2016**;7:11500-11
10. Molina JR, Sun YT, Protopopova M, Gera S, Bandi M, Bristow C*, et al.* An inhibitor of oxidative phosphorylation exploits cancer vulnerability. Nat Med **2018**;24:1036-46
11. Souers AJ, Leverson JD, Boghaert ER, Ackler SL, Catron ND, Chen J*, et al.* ABT-199, a potent and selective BCL-2 inhibitor, achieves antitumor activity while sparing platelets. Nat Med **2013**;19:202-8
12. Kotschy A, Szlavik Z, Murray J, Davidson J, Maragno AL, Le Toumelin-Braizat G*, et al.* The MCL1 inhibitor S63845 is tolerable and effective in diverse cancer models. Nature **2016**;538:477-82
13. Donati G, Peddigari S, Mercer CA, Thomas G. 5S ribosomal RNA is an essential component of a nascent ribosomal precursor complex that regulates the Hdm2-p53 checkpoint. Cell Rep **2013**;4:87-98
14. Sanchez-Arevalo Lobo VJ, Doni M, Verrecchia A, Sanulli S, Faga G, Piontini A*, et al.* Dual regulation of Myc by Abl. Oncogene **2013**;32:5261-71
15. D'Andrea A, Gritti I, Nicoli P, Giorgio M, Doni M, Conti A*, et al.* The mitochondrial translation machinery as a therapeutic target in Myc-driven lymphomas. Oncotarget **2016**;7:72415-30
16. Campos CB, Paim BA, Cosso RG, Castilho RF, Rottenberg H, Vercesi AE. Method for monitoring of mitochondrial cytochrome c release during cell death: Immunodetection of cytochrome c by flow cytometry after selective permeabilization of the plasma membrane. Cytometry A **2006**;69:515-23
17. Bianchi V, Ceol A, Ogier AG, de Pretis S, Galeota E, Kishore K*, et al.* Integrated Systems for NGS Data Management and Analysis: Open Issues and Available Solutions. Front Genet **2016**;7:75
18. Gentleman RC, Carey VJ, Bates DM, Bolstad B, Dettling M, Dudoit S*, et al.* Bioconductor: open software development for computational biology and bioinformatics. Genome Biol **2004**;5:R80
19. Love MI, Huber W, Anders S. Moderated estimation of fold change and dispersion for RNA-seq data with DESeq2. Genome Biol **2014**;15:550
20. Subramanian A, Tamayo P, Mootha VK, Mukherjee S, Ebert BL, Gillette MA*, et al.* Gene set enrichment analysis: a knowledge-based approach for interpreting genome-wide expression profiles. Proc Natl Acad Sci U S A **2005**;102:15545-50
21. Liberzon A, Birger C, Thorvaldsdottir H, Ghandi M, Mesirov JP, Tamayo P. The Molecular Signatures Database (MSigDB) hallmark gene set collection. Cell Syst **2015**;1:417-25
22. Monti S, Savage KJ, Kutok JL, Feuerhake F, Kurtin P, Mihm M*, et al.* Molecular profiling of diffuse large B-cell lymphoma identifies robust subtypes including one characterized by host inflammatory response. Blood **2005**;105:1851-61
23. Edgar R, Domrachev M, Lash AE. Gene Expression Omnibus: NCBI gene expression and hybridization array data repository. Nucleic Acids Res **2002**;30:207-10
24. Ran FA, Hsu PD, Wright J, Agarwala V, Scott DA, Zhang F. Genome engineering using the CRISPR-Cas9 system. Nat Protoc **2013**;8:2281-308
25. Ianevski A, He L, Aittokallio T, Tang J. SynergyFinder: a web application for analyzing drug combination dose-response matrix data. Bioinformatics **2017**;33:2413-5
26. Lenz G, Wright GW, Emre NC, Kohlhammer H, Dave SS, Davis RE*, et al.* Molecular subtypes of diffuse large B-cell lymphoma arise by distinct genetic pathways. Proc Natl Acad Sci U S A **2008**;105:13520-5
27. Reddy A, Zhang J, Davis NS, Moffitt AB, Love CL, Waldrop A*, et al.* Genetic and Functional Drivers of Diffuse Large B Cell Lymphoma. Cell **2017**;171:481-94 e15
28. Chapuy B, Stewart C, Dunford AJ, Kim J, Kamburov A, Redd RA*, et al.* Molecular subtypes of diffuse large B cell lymphoma are associated with distinct pathogenic mechanisms and outcomes. Nat Med **2018**;24:679-90
29. Schmitz R, Wright GW, Huang DW, Johnson CA, Phelan JD, Wang JQ*, et al.* Genetics and Pathogenesis of Diffuse Large B-Cell Lymphoma. N Engl J Med **2018**;378:1396-407
30. Ennishi D, Jiang A, Boyle M, Collinge B, Grande BM, Ben-Neriah S*, et al.* Double-Hit Gene Expression Signature Defines a Distinct Subgroup of Germinal Center B-Cell-Like Diffuse Large B-Cell Lymphoma. J Clin Oncol **2019**;37:190-201
31. Sha C, Barrans S, Cucco F, Bentley MA, Care MA, Cummin T*, et al.* Molecular High-Grade B-Cell Lymphoma: Defining a Poor-Risk Group That Requires Different Approaches to Therapy. J Clin Oncol **2019**;37:202-12

**Supplemental Table 1**: Genes shared among the MYC- and OxPhos-related signatures

| Gene set 1 (nr. of genes) | Gene set 2 (nr. of genes) | Nr. of overlapping genes |
|---|---|---|
| Hallmark-OxPhos (200) | MYC-V1 (200) | 11 |
| Hallmark-OxPhos (200) | MYC-V2 (58) | 1 |
| Hallmark-OxPhos (200) | CCC-OxPhos (56) | 18 |
| CCC-OxPhos (56) | MYC-V1 (200) | 3 |
| CCC-OxPhos (56) | MYC-V2 (58) | 0 |
| MYC-V1 (200) | MYC-V2 (58) | 18 |

**Supplemental Table 2**: Antibodies used for immunoblot analysis

| Target protein | Source | Clone | Vendor | Cat # |
|---|---|---|---|---|
| Vinculin | mouse | hVIN-1 | Merck Life Science | V9131 |
| Myc | mouse | Y69 | Abcam | ab32072 |
| PARP | rabbit | polyclonal | Cell Signaling Technology | 9542 |
| BCL2 | rabbit | D17C4 | Cell Signaling Technology | 3498 |
| Bcl-XL | rabbit | E18 | Abcam | ab32370 |
| Bax | mouse | E63 | Merck Life Science | ab32503 |
| Bim | mouse | C34C5 | Cell Signaling Technology | 2933 |
| PUMA a/b | rabbit | polyclonal | Santa Cruz Biotechnologies | sc-28226 |
| Phospho-eIF2a (S51) | rabbit | D9G8 | Cell Signaling Technology | 3398 |
| eIF2a | rabbit | D7D3 | Cell Signaling Technology | 5324 |
| ATF4 | rabbit | D4B8 | Cell Signaling Technology | 11815 |
| CHOP | mouse | L63F7 | Cell Signaling Technology | 2895 |
| Pck2 | rabbit | polyclonal | Cell Signaling Technology | 6924 |
| Gpx1 | rabbit | polyclonal | Abcam | ab22604 |

**Supplemental Table 3**: Primers for mRNA quantification by qPCR

| Target gene | NCBI gene ID | Forward primer sequence | Reverse primer sequence |
|---|---|---|---|
| Tbp | 21374 | TCAAACCCAGAATTGTTCTCC | TTCAAATGCTTCATAAATCTCTGC |
| Myc | 17869 | TTTTTGTCTATTTGGGGACAGTG | CATCGTCGTGGCTGTCTG |
| St6Galnac4 | 20448 | TGGTCTACGGGATGGTCA | CTGCTCATGCAAACGGTACAT |
| Rrp9 | 27966 | TTCTAGCGGACGCGATAAAC | ACTTCTGCAACCTGCCTCTC |

**Supplemental Table 4**: genomic targets of the sgRNAs used for CRISPR-Cas9 gene knockout

| Target gene | NCBI gene ID | sgRNA target sequence | Locus |
|---|---|---|---|
| Ddit3 | 13198 | TCAGCTGCCATGACTGCACG | chr10:127295332-127295351 |
| Ddit3 | 13198 | CTGTCCTCAGATGAAATTGG | chr10:127295425-127295444 |
| Bax | 12028 | AGCGAGTGTCTCCGGCGAAT | chr7:45466013-45466032 |
| Bax | 12028 | AGTTTCATCCAGGATCGAGC | chr7:45466106-45466125 |
| Bak | 12018 | TCATCGCAGCCCACCTTCGG | chr17:27025810-27025829 |
| Bak | 12018 | CTGGTGTGCGCACATGCGCA | chr17:27025715-27025734 |

**Supplemental Table 5**: Primers for PCR analysis of CRISPR-Cas9 knockout clones

| Target gene | NCBI gene ID | Forward primer sequence | Reverse primer sequence | Amplicon |
|---|---|---|---|---|
| Ddit3 | 13198 | CCCATGCCCTTACCTATCGT | AGGAGAGGCATACAAACCCC | chr10:127295211-127295680 |
| Bax | 12028 | AACATTCTGCTCCTCTCCCC | CAGAGCACCGCCTACTAGAA | chr7:45465728-45466162 |
| Bak | 12018 | AGGTCACACATCACTACCCG | AATGCCATTCCCTGTCCACA | chr17:27025495-27025949 |

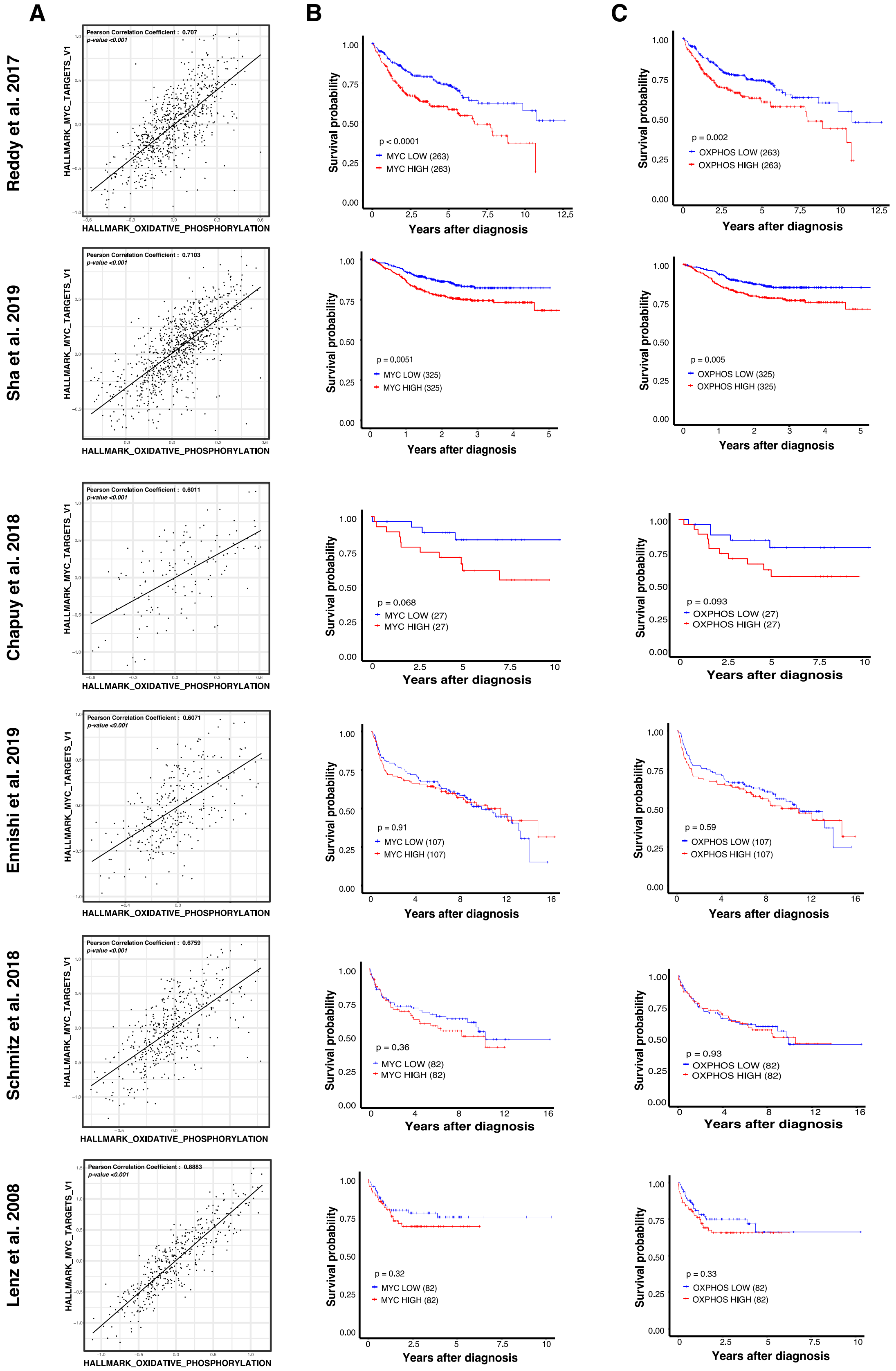


**Supplemental Figure 1. MYC- and OxPhos-associated gene signatures are correlated in DLBCL.** Gene expression profiles from 6 independent human DLBCL patient cohorts (26-31) (left; as in Table 1) were used in our analyses. (**A**) Correlation between the Hallmark OxPhos and MYC-V1 gene sets. The X and Y axes report the mean expression of all genes in the indicated signature, in each patient (dots). Black lines represent linear regression fits to the data points. (**B, C**) Kaplan-Meier survival curves for R-CHOP treated DLBCL patients stratified according to the expression of genes in Hallmark MYC-V1 (B) or OxPhos (C); the plots compare the survival of patients grouped in the top (HIGH) and bottom (LOW) tertiles, with the numbers of patients in each group given in parenthesis. Note all cases with transcriptional data were considered in (A), but only those with associated clinical information in (B, C). For the study of Lenz et al. (26), only R-CHOP-treated patients were analyzed, excluding different treatments; in Sha et al. (31) (REMoDL-B trial), about 50% of the patients were treated with bortezomib in addition to R-CHOP, and were included in our analyses.

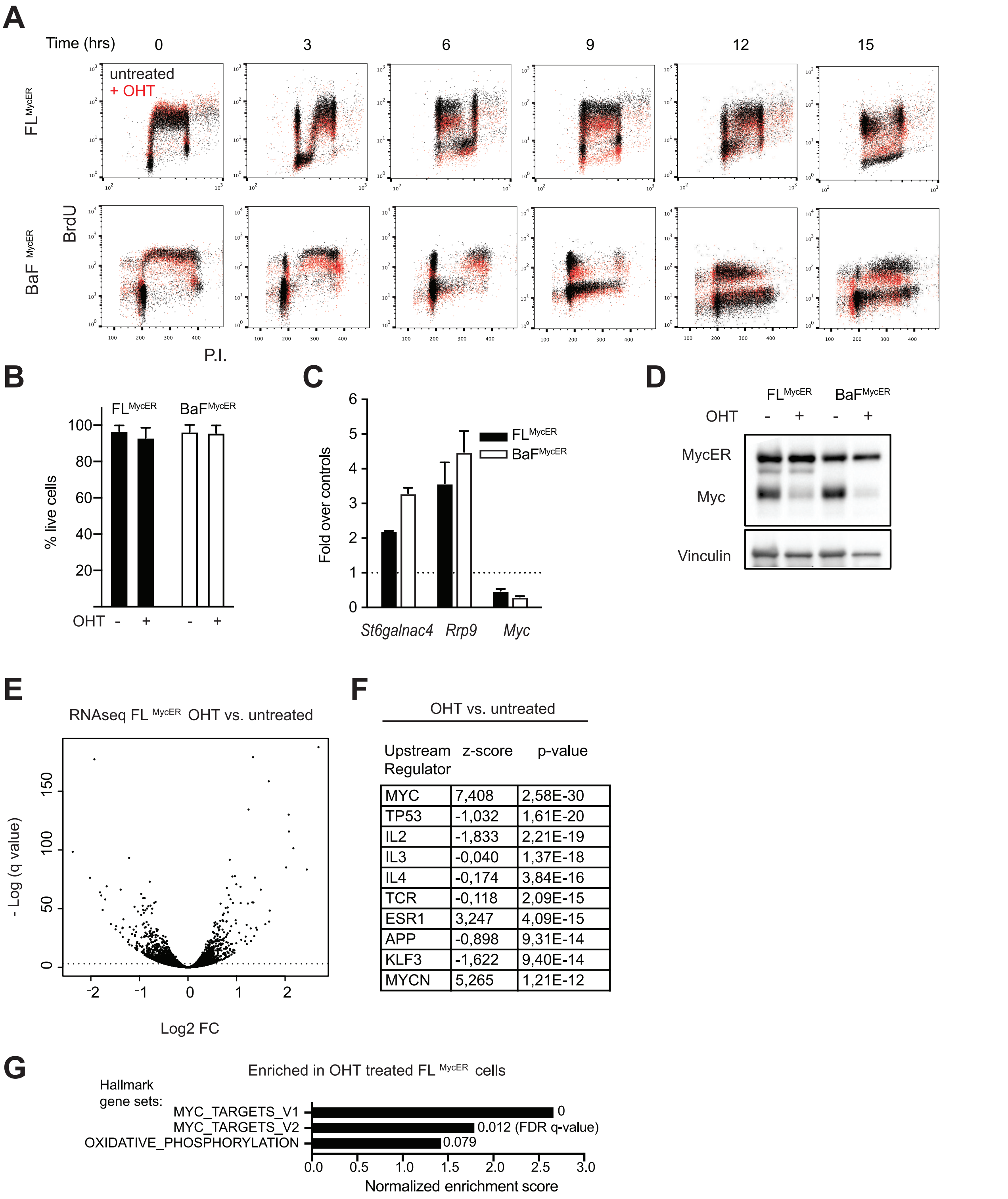


| Upstream Regulator | z-score | p-value |
|---|---|---|
| MYC | 7,408 | 2,58E-30 |
| TP53 | -1,032 | 1,61E-20 |
| IL2 | -1,833 | 2,21E-19 |
| IL3 | -0,040 | 1,37E-18 |
| IL4 | -0,174 | 3,84E-16 |
| TCR | -0,118 | 2,09E-15 |
| ESR1 | 3,247 | 4,09E-15 |
| APP | -0,898 | 9,31E-14 |
| KLF3 | -1,622 | 9,40E-14 |
| MYCN | 5,265 | 1,21E-12 |

**Supplemental Figure 2. MycER expression and activation in the B-lymphoid cell lines FL5.12 and Ba/F3.** Following transduction with a MycER-expressing retrovirus and selection of resistant cell pools ($FL^{MycER}$ and $BaF^{MycER}$), the constitutively expressed MycER chimera was activated by treatment with OHT (100 nM, 48h). (**A**) Cell cycle kinetics: cells were pulse-labeled with BrdU (33 µM, 20 minutes), chased in BrdU-free medium and harvested at the indicated time points. The profiles of OHT-treated and untreated cells (red and black dots, respectively) are overlaid, revealing undistinguishable progression kinetics. (**B**) Percentage of live cells following 48h OHT treatment. Error bars: SD (n=3). (**C**) RT-PCR quantification of endogenous *MYC* and known MYC-induced mRNAs, normalized to *Tbp*. (**D**) Immunoblot analysis with a MYC-specific antibody, detecting both endogenous mouse MYC and exogenous MycER, as indicated. Vinculin was used as loading control. (**E**) Volcano plot showing fold change (FC, Log2 value) against q-value (–log10) for each mRNA in OHT-treated (100 nM, 72h) vs. untreated $FL^{MycER}$ cells. The threshold of statistical significance for calling Differentially Expressed Genes (DEGs; q value < 0.05) is identified by the dotted line. (**F**) 10 most significantly enriched upstream regulators identified from OHT-responsive DEGs. (**G**) Enrichment of MYC target and OxPhos gene sets among OHT-responsive DEGs.

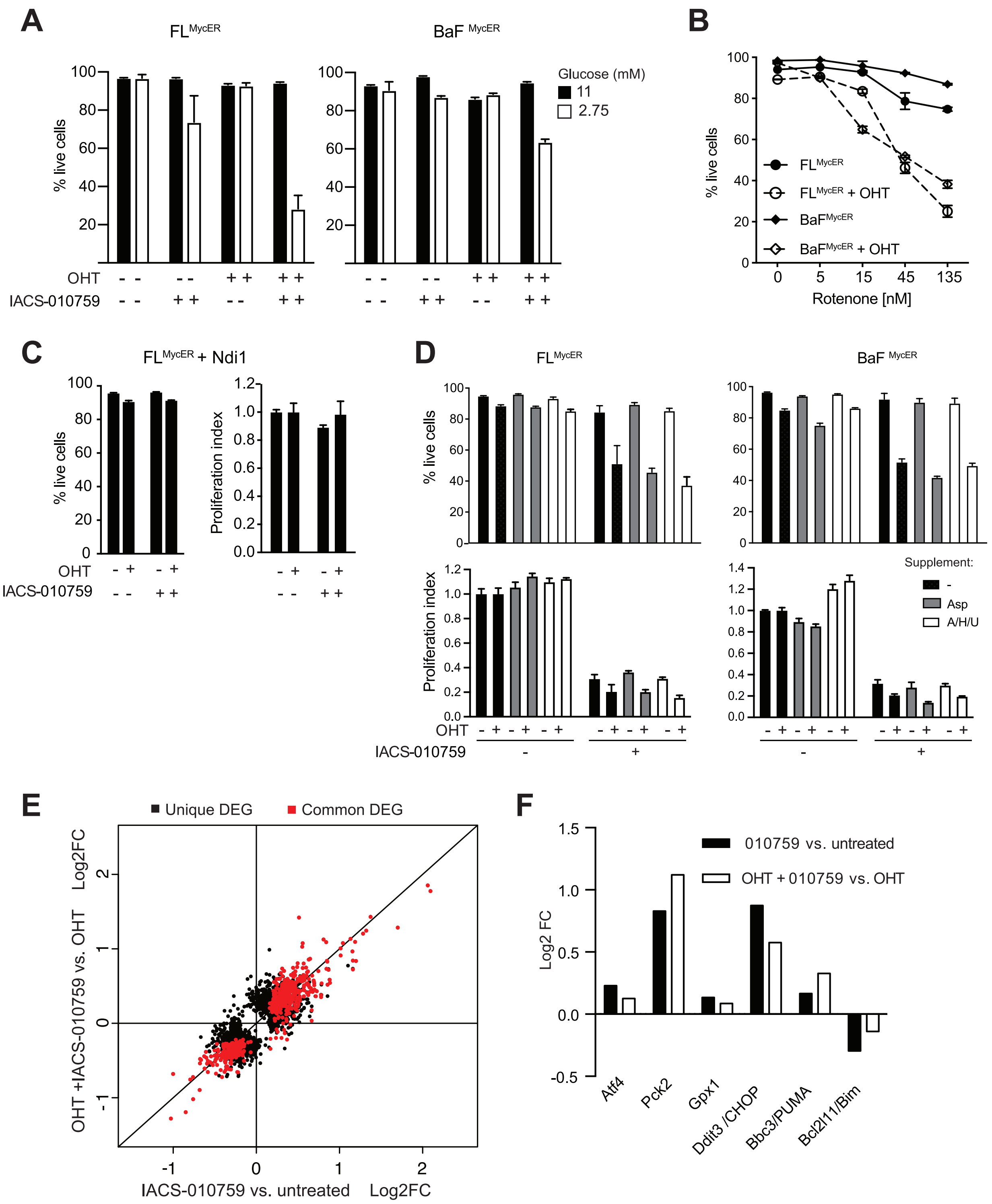


**Supplemental Figure 3. Effects of MycER activation, IACS-010759 and other pharmaco-genetic interactions in B-cells.** (**A**) Percentage of live $FL^{MycER}$ and $BaF^{MycER}$ cells following sequential treatment with OHT (100 nM, 48h) and IACS-010759 (135 nM, 48h) at the indicated glucose concentrations. (**B**) As in (A), following sequential treatment with OHT and rotenone (48h) at the indicated concentrations. (**C**) Percentage of live cells (left) and Proliferation index (right; as defined in Fig. 2A) following the indicated treatments in Ndi1-expressing $FL^{MycER}$ cells. (**D**) Same as (C) in $FL^{MycER}$ and $BaF^{MycER}$ cells treated as indicated with OHT and IACS-010759 (135 nM) for 48h in standard growth medium, or medium supplemented with either aspartate (Asp; 10 mM), or a combination (A/H/U) of adenine (150 µM), hypoxantine (150 µM), and uridine (400 µM). Error bars in A-D: SD (n=3). (**E**) RNA-seq profiling of the response to IACS-010759 (135 nM, 24 hours), either with (Y-axis) or without (X-axis) pre-treatment with OHT (100 nM, 48h). Black dots mark transcripts called as DEGs (q value < 0.05) in only one condition, and red dots those called in both. (**F**) Variation of known ISR-regulated mRNAs upon IACS-010759 treatment in cells grown with or without OHT from the RNA-seq data.

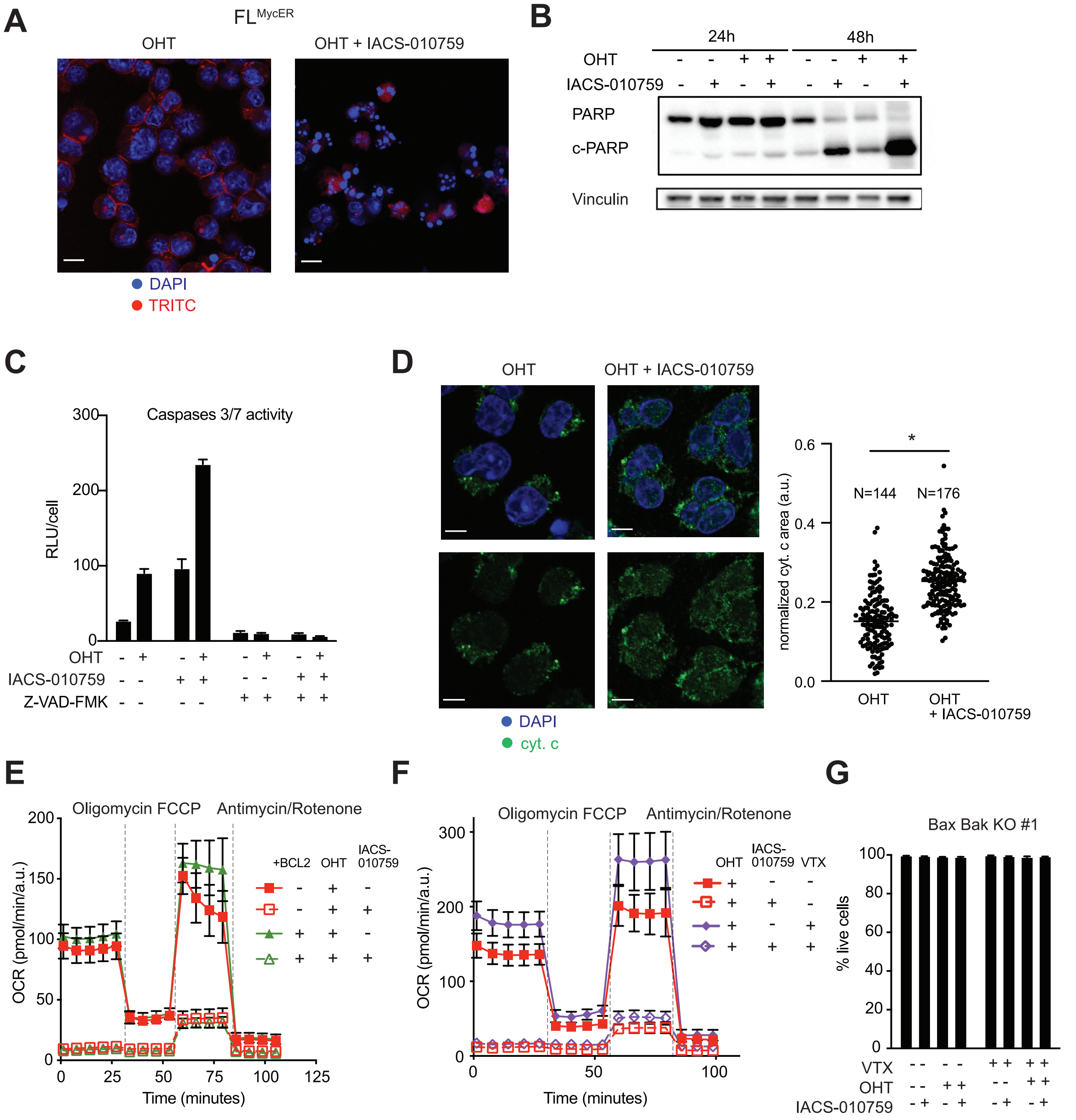


**Supplemental Figure 4. IACS-010759 kills FL$^{MycER}$ cells by apoptosis.** FL$^{MycER}$ cells were primed with OHT and treated with IACS-010759 (135 nM, 48h, unless otherwise indicated). (**A**) DAPI and TRITC-Agglutinin staining. In the IACS-010759-treated sample, note the presence of nuclei with condensed and fragmented chromatin, or faintly stained due to DNA loss. Scale bars: 10 µm. (**B**) Immunoblot analysis showing the cleaved (c-PARP) and uncleaved forms of PARP. Vinculin was used as loading control. (**C**) Caspase 3/7 activity in FL$^{MycER}$ cells treated with OHT, IACS-010759 and/or 20 µM Z-VAD-FMK (48h) as indicated. (**D**) Representative images of DAPI and cytochrome c staining (left) and relative quantification of the cytochrome c cellular area (right). *P < 0.0001 (Student's t test). Scale bars: 10 µm. (**E, F**) Mitochondrial stress test profiles of control and BCL2-overexpressing FL$^{MycER}$ cells, primed with OHT and treated with IACS-010759 (135 nM) and/or venetoclax (100 nM) for 24 hours, as indicated. (**G**) Percentage of live Bax/Bak knockout FL$^{MycER}$ cells following treatment with IACS-010759 (135 nM) and/or venetoclax (100 nM), as indicated. Error bars: SD (n=3).

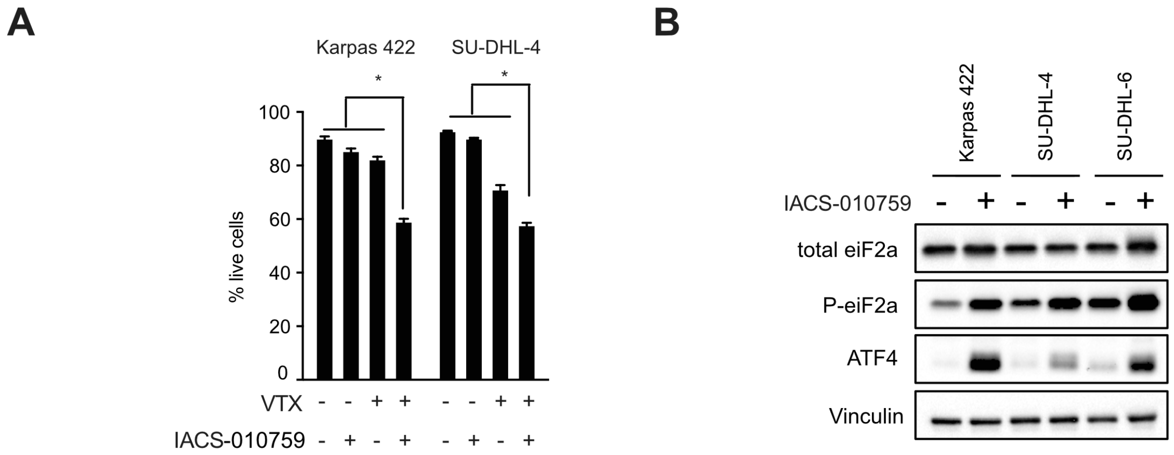


C

| | Compound | Dose (mg/kg) | Partial Regression (Day 19) | Complete Regression (Day 19) | Tumor free (Day 120) |
|---|---|---|---|---|---|
| Experiment 1 | VTX | 50 | 0/5 | 0/5 | 0/5 |
| | IACS-010759 | 1 | 0/5 | 0/5 | 0/5 |
| | IACS-010759 | 5 | 0/5 | 0/5 | 0/5 |
| | IACS-010759 | 10 | 0/5 | 0/5 | 0/5 |
| | IACS-010759 | 15 | 1/4 | 0/4 | 0/4 |
| | VTX + IACS | 50 + 1 | 0/5 | 0/5 | 0/5 |
| | VTX + IACS | 50 + 5 | 3/5 | 1/5 | 1/5 |
| | VTX + IACS | 50 + 10 | 3/5 | 2/5 | 0/5 |
| | VTX + IACS | 50 +15 | 1/4 | 3/4 | 0/4 |
| Experiment 2 | VTX | 50 | 0/5 | 0/5 | 0/5 |
| | IACS-010759 | 2,5 | 0/5 | 0/5 | 0/5 |
| | IACS-010759 | 5 | 0/5 | 0/5 | 0/5 |
| | VTX + IACS | 50 + 2,5 | 3/5 | 1/5 | 0/5 |
| | VTX + IACS | 50 + 5 | 3/5 | 2/5 | 1/5 |

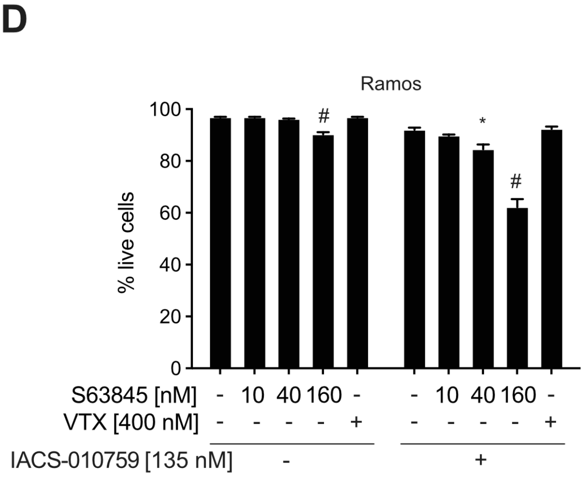


**Supplemental Figure 5. Effects of IACS-010759 and BH3-mimetics on human lymphoma cells.** (**A**) Percentage of live cells after treatment (24h) of the human DHL cell lines Karpas 422 and SU-DHL-4 with IACS-010759 (135 nM) and/or venetoclax (VTX, 100 nM), as indicated. *P < 0.0001. (**B**) Immunoblot analysis of total, phosphorylated (P-) eIF2a and ATF4 in the indicated DHL cell lines, treated with IACS-010759 (135 nM) for 24 hours. Vinculin was used as loading control. (**C**) Summary of the response of subcutaneous SU-DHL-6 tumors to treatment with IACS-010759 and/or venetoclax, as indicated, in two independent experiments. Detailed progression curves for experiment 1 are shown in Figure 5C. Partial and Complete Regression at day 19 (one week after the end of treatment) were defined as ≤50% and ≤10% of the starting tumor volume (day 0), respectively. (**D**) The Burkitt's lymphoma cell line Ramos was treated for 24h with IACS-010759 (135 nM), alone or in combination with venetoclax or S63845, as indicated. The plot reports the percentage of live cells after treatment. *P < 0.01 and #P < 0.0001vs. the appropriate control (untreated or treated with IACS-010759 only). Error bars: SD (n=3).
